# Measuring peptide–MHC generalization to unseen alleles across both HLA classes

**DOI:** 10.64898/2026.06.18.733075

**Authors:** Venkatesh Mysore

## Abstract

Reported peptide–MHC (pMHC) AUROCs of 0.85–0.95 overstate generalization to unseen alleles: because immunopeptidome data are dense on a few well-studied alleles and sparse on the rest, training and test sets come to share near-identical alleles, so the numbers partly reflect interpolation rather than extrapolation to new MHC grooves. This is a property of the data, not of any one method. We assembled an open, harmonized corpus of 5.8 million experimental measurements across both HLA classes and use it to control the leakage explicitly: alleles held out at the sequence and cluster level, peptide-disjoint splits, and provenance-matched negatives. On strictly novel alleles, generalization is in the high 0.7s rather than the 0.9s a conventional split returns. Against this benchmark we trained a predictor that spans both classes in one model and factors presentation into a peptide-only ligand-likeness term and an allele-specific term; it exceeds eight published predictors by per-allele Δ_AUROC_ = +0.22 to +0.37 (*p <* 10*^−^*^9^), most on the least-studied genes. Corpus, benchmark, and model are released.

**Author summary:** Our immune cells display protein fragments on the cell surface, held by molecules (the human leukocyte antigens, or HLAs) that vary from person to person. Predicting which fragments a given HLA displays matters for cancer vaccines, transplant matching, and the safety of engineered therapies, and many computational tools now do it well. Most available data come from a few common HLAs, so test cases tend to resemble training cases, and the published accuracy looks better than it really is for the rare HLAs that matter most in the clinic. We assembled a large, openly shared collection of experimental measurements across both major HLA classes and used it to test prediction more directly, holding out HLAs that are sequence-distant from those in training. Accuracy on these is measurable but lower than the usual figures suggest. We also built a predictor that handles both HLA classes in one model and gains most relative to existing tools on the rare HLAs where they are weakest. The data, benchmark, and model are available for the same test.

## Introduction

Antigen presentation by major histocompatibility complex (MHC) molecules is central to adaptive immunity, and predicting which peptides bind which MHC alleles is foundational to the de-immunogenization of protein and peptide therapeutics, personalised cancer vaccines ^14,33^, transplant compatibility algorithms^1^, TCR-mimetic and adoptive-cell safety screens^25,34^, and pMHC-based tolerogenic nanomedicines^2^. These applications span both classes of human leukocyte antigen (HLA, the human MHC) and need calibrated probabilities rather than rank percentiles^20^. The NetMHCpan family ^3,21^ and an active open-source community ^5,6,12,13,17,18^ report headline AUROCs of 0.85–0.95 on their held-out partitions.

Measuring generalization to previously unseen alleles or peptides is intrinsically hard, because pMHC data accrete by re-deposition of shared peptides and alleles across studies. Two features of current practice can therefore yield an over-optimistic estimate. First, training and test sets overlap at the sequence level. The standard “novel-allele” split holds out alleles by name but can leave training alleles at very high identity to test examples ^27,28^. In our audit, 73–83% of test alleles in such splits had a training neighbor at ≥0.95 G-domain identity, so the resulting AUROCs largely reflect interpolation between near-identical training alleles rather than extrapolation to unseen grooves ^18,29^. This overlap is rarely measured per tool. Second, generated negatives can carry a distributional signature that even a peptide-only classifier can detect, allowing a model to gain AUROC from provenance rather than binding. These leakage modes parallel the benchmark-contamination concerns of machine learning more broadly^11,19,35,40^; structure-based work has already shown, for Class I, that allele-clustered splits reveal this gap ^18^, and we build on that line by extending it to a pan-class, sequence-level setting with the negative-generation and audit checks a trainable corpus requires.

We use a leakage-controlled benchmark that pairs a disjoint hold-out with matched negatives. The held-out stratum is disjoint on the peptide and the MHC axis at once: every test allele lies in a different 90%-identity cluster from every training allele, and below 0.90 G-domain identity to all of them, while every test peptide lies in a different 80%-identity cluster from every training peptide. A measured allele is then sequence-distant rather than a renamed near-twin. The negatives are source-matched: each decoy is a binder to a different allele from the same source, so a peptide-only baseline cannot tell decoys from real negatives by provenance. We release the contamination audit that checks these properties, so the benchmark can be reused on a rebuilt or extended corpus. For each comparator, we also remove test rows present in that tool’s training data.

We also release Aiki-HLA, a class-agnostic predictor that feeds unmodified ESM-2 650M embeddings^4,16,26^ of the full MHC G-domain to a single attention head. The enabling choice is a common 182-residue input for both classes: the Class I single-chain *α*1+*α*2 groove, and the Class II *α*1+*β*1 heterodimer flattened into one length-matched sequence ^36^ rather than modeled as a dimer. This deliberate simplification uses one set of weights for both classes, whereas prior tools use a 34-residue pseudosequence^27,28^ and class-specific models ^37,38^. We adopt the full G-domain for what it enables, a single class-agnostic head, not for accuracy: on our strict stratum it is statistically equivalent to the pseudosequence (SI Table 5).

Aiki-HLA is released as two ensembles of identical architecture and training recipe that differ only in the licensing of their source data: an **open ensemble** on a fully-redistributable, commercial-use-permitted subset (classical human HLA), and a **research-use ensemble** that adds non-commercial sources, non-classical and MHC class I-related human loci, and mass-spectrometry-derived allele assignments (corpus sizes in Results). The corpus, embeddings, contamination audit, and both ensembles are released with the benchmark.

Aiki-HLA reflects two design choices. One is to score both HLA classes with a single model over a common representation, rather than the separate Class I and Class II models the field has usually built. The other is to factor presentation into the two questions implicit in the prediction task: is this peptide an MHC ligand at all, and, if so, does it fit this particular allele’s groove? An allele-blind *ligand-likeness* score *p*_ligand_ estimates the first from peptide features alone (no allele input), Aiki-HLA’s allele-specific score *p*_binding_ estimates the second, and the score we release for downstream use is their geometric mean 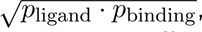, a calibrated presentation probability with no fitted mixing parameter. Each factor addresses a different part of the task: general ligand recognition, where existing tools already do well, and allele specificity, where rare alleles remain difficult. Their product is therefore usable in both settings, and *p*_binding_ remains available where the task is itself allele-specific (cross-tool comparison, novel-allele discrimination).

## Results

### Training corpora and model ensembles

The **open ensemble** is trained on 4,662,506 experimental measurements over 421 classical HLA-A/B/C/DR/DP/DQ alleles, from sources whose licenses permit unrestricted redistribution: the partition on which every comparator can be scored without a licensing confound. The **research-use ensemble** extends it to 5,796,339 measurements over 507 alleles by adding non-commercial sources, non-classical HLA, and deconvolution-derived assignments (Methods). Releasing the pair separates two effects a single larger model would confound: *coverage broadening* (111 alleles the open corpus cannot score, taking the evaluable set from 174 to 285) and better performance on the 174 that both cover. Both use source-matched negatives, which keep provenance from leaking into the label (Methods).

Distinct from both is a third, *evaluation ensemble*: the model used to define the benchmark. Trained on a subset of the open corpus from which every training allele within a 0.85–0.92 identity band of a held-out test allele is removed (a bridge purge, Methods), it evaluates extrapolation to a sequence-distant groove rather than interpolation from a near-twin. The 0.765 strict lower bound below is therefore a measurement of novel-allele generalization rather than near-neighbor interpolation. It is a benchmark model, not a deployed predictor; its splits and purged corpus are released so any pMHC model can be placed on the same footing.

The allele-blind ligand-likeness score *p*_ligand_ is trained separately, on presented peptides alone: 1.70 million distinct eluted ligands (the positive peptides of both corpora, pooled across alleles) against length-matched human-proteome fragments, with no allele input. The recent-IEDB peptides held out for testing are excluded from its training too, so the presentation score is evaluated on peptides neither factor has seen (Methods).

### Generalization to novel alleles

Because performance depends on how far a query allele sits from the training set, we report a two-sided bracket. The *lower bound* measures only strictly novel alleles, held out from every training allele at both the cluster and the raw-sequence level, with peptide-disjoint splits (Methods); the *upper bound* measures a conventional held-out set not controlled for sequence novelty, the setting most published tools report. A query near the training set performs near the upper bound; a query with no close relative falls toward the lower one.

#### Lower bound

On the 39 strictly novel alleles (maximum G-domain identity to any training allele below 0.90), restricted to the 140,073 directly measured rows on those alleles (real assayed non-binders, no generated decoys), the evaluation ensemble reaches a per-allele median AUROC of **0.765** [0.691, 0.797] (Class I 0.765 on 23 alleles; Class II 0.709 on 16; Table 1). Because the two released ensembles train on more data, 0.765 is a conservative floor for both. A peptide-only logistic regression on the same alleles reaches only **0.493** [0.447, 0.527] (peptide length alone 0.509), so the **0.272**-point gain comes from reading the MHC groove, not from peptide sequence alone (Fig. S1b). These 39 alleles fall into three novel G-domain neighborhoods: one HLA-C group (23 variants, nearest training neighbor an HLA-B allele at 0.879–0.896 identity) and two HLA-DQ heterodimer groups. Because the result reproduces across the natural variants within each neighborhood, the 0.765 reflects extrapolation to unseen grooves, not an artifact of any single allele. Three stricter tests appear in Supplementary Tables 4 and 8: 16 HLA-DQ heterodimers at 0.802–0.835 identity, a 0.90–0.93 marginal band at PA-med 0.661, and an automatic gene-family bridge purge that lowers PA-med by ~0.12. Test hardness is allele-driven, not peptide-driven (Supplementary Fig. S3a). MHCflurry-2.0 degrades along the same allele-distance axis (Supplementary Fig. S3b), which places the frontier in the training data rather than in any one model. **Upper bound.** On a conventional held-out set, the open ensemble reaches a per-allele median AUROC of **0.913** [0.904, 0.924] (70,781 rows withheld from training, stratified by class and source; Class I 0.923, Class II 0.900). The research-use ensemble reaches a statistically equivalent **0.911** [0.893, 0.927] while covering 64% more alleles (285 evaluable, 441,443 rows). The two ensembles agree closely on the 174 alleles both can evaluate (research-use 0.885, Δ = −0.018, traced mostly to a train/test negative-generation mismatch; Methods). The added data instead extends coverage to 111 rare subtypes the redistributable corpus could not support at all, and on those it reaches **0.949**. This upper bound falls in the 0.85–0.95 range published tools report^3,12^, as expected for a conventional split. User-visible performance therefore lies in the bracket [0.71, 0.92], set by how novel the query allele is.

**Table 1:**
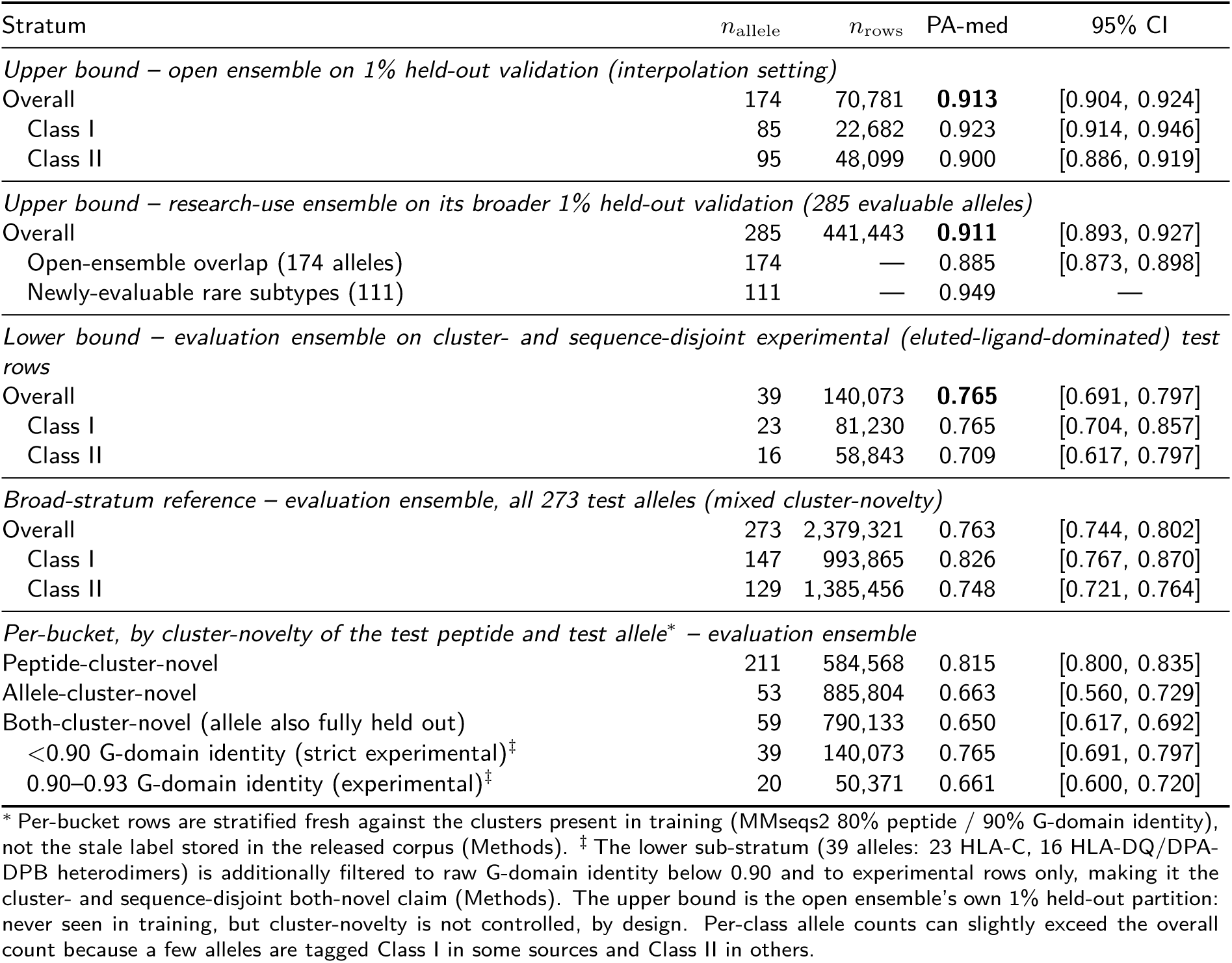
Aiki-HLA performance bracket on cluster-novel (lower bound) and interpolation (upper bound) settings. User-visible AUROC for a given query lies between the lower-bound and upper-bound rows depending on how cluster-novel the query is against the training corpus. The *lower bound* comes from a bridge-purged eval ensemble (3 seeds, recipe-identical to the open ensemble, trained on a corpus from which within-family training alleles within 0.85–0.92 G-domain identity to each strict-stratum test allele are removed before training) scored on the cluster- and sequence-disjoint sub-stratum. The open-subset *upper bound* comes from the open ensemble on its own 1% held-out validation partition (stratified by class and source; the conventional random-stratified held-out protocol used by most published pMHC tools). The broader *upper bound* comes from the research-use ensemble on its own 1% held-out validation partition; of the 285 evaluable alleles, 174 overlap the open ensemble’s evaluable set (research-use 0.885 [0.873, 0.898]; pure-corpus Δ = 0.018, 0.011 residual after the experimental-only control for train/test decoy-method mismatch; Methods), and the remaining 111 are newly-evaluable rare-subtype alleles. All values are per-allele median AUROC; CIs are 1000-sample bootstraps over alleles within each stratum.

Across the broad 273-allele test stratum, difficulty increases with novelty as the bracket implies (novel-peptide 0.815 *>* novel-allele 0.663 *>* both-novel 0.650; Table 1), and per-gene performance tracks data density (HLA-B 0.896 strongest, HLA-C 0.659 weakest). Calibration is class-asymmetric (Brier 0.140; expected calibration error 0.15 Class I, 0.05 Class II) and degrades with novelty (Supplementary Fig. S2). Across seeds the result is stable (0.745 ± 0.011).

### Head-to-head against eight existing tools

The comparison uses rows absent from each tool’s training data: for each comparator we remove the (peptide, allele) pairs in its own training data before scoring (Methods). On the open ensemble’s 1% held-out validation partition we evaluate eight published predictor variants^3,5,6,12,13,28^, each on its supported human-HLA alleles, with no species or licensing confound (Table 2).

**Table 2:**
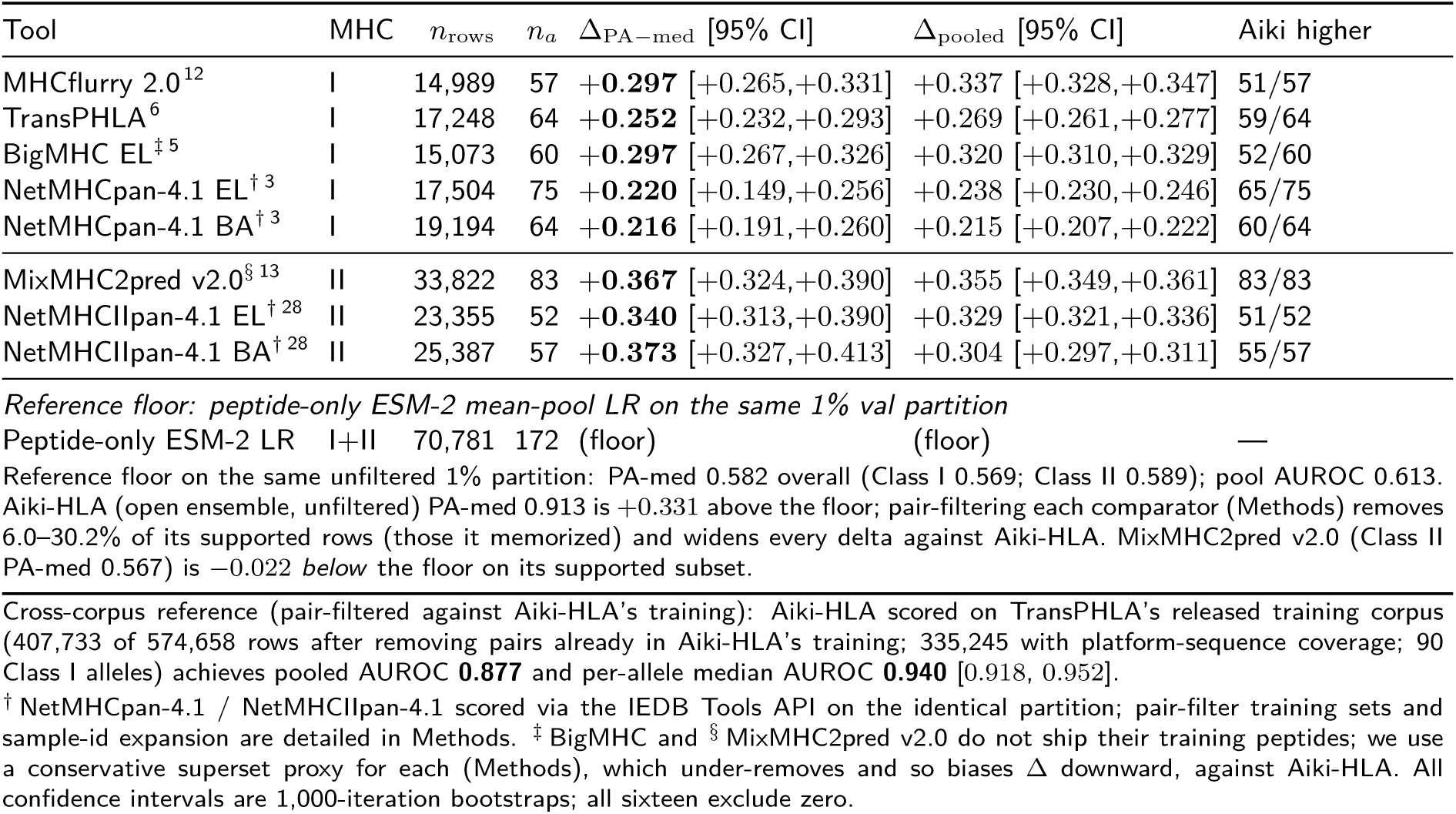
Cross-tool head-to-head on the open ensemble’s 1% held-out validation partition (pair-symmetric protocol). The protocol measures allele-specific discrimination within the eluted-ligand manifold (positives versus source-matched cross-allele negatives); it is distinct from the common-allele proteome-decoy screening task existing tools are built for, on which they match or exceed Aiki-HLA (Discussion, “Presentation versus affinity”). The 1% partition (70,781 rows, stratified by mhc_class source) was held out from the open ensemble’s training. For every comparator we additionally remove from the test subset any exact (peptide, allele) pair that appears in that comparator’s training data, so memorization cannot carry the comparison (sample-id expansion, and the MixMHC2pred-v2.0 proxy training set marked , are detailed in Methods). *n*_rows_ is the paired-prediction row count after the comparator’s allele + length filters *and* the pair-level filter; *n*_allele_ is the subset with ≥30 paired rows and both classes present. Δ_PA−med_ is the per-allele-median paired AUROC of the open Aiki-HLA ensemble minus the comparator; Δ_pooled_ is the row-pooled AUROC difference. Both metrics agree in sign across all eight comparator flavors; Wilcoxon *p <* 10^−9^ on every row.

The reference predictor has higher per-allele median AUROC by Δ_AUROC_ = +0.22 to +0.37, winning a majority of alleles against each (52 to 83 alleles, 15,000–34,000 paired rows per comparator; Wilcoxon *p <* 10^−9^), with the row-pooled difference agreeing in sign in every case. On Class I alleles with ≥100 paired rows after pair-filter (*n_a_* = 40 to 43 depending on comparator), Aiki-HLA places real binders in its top 100 with 0.83 to 0.89 precision against 0.44 to 0.61 for the three Class I comparators on the same protocol, the shortlist-relevant metric for applications that examine only the head of a ranking. The margin is largest on the least-studied genes (HLA-C and the HLA-DQ heterodimers), consistent with a benchmark built to measure generalization to under-represented alleles. Two qualifiers temper this. The margin is in allele-specific discrimination, not bulk-proteome screening: on a companion proteome-decoy task the predictor and the strongest comparator are statistically tied (Δ_AUROC_ = − 0.011; Discussion). And the training corpus is 5–10× larger than every comparator’s, so part of the margin reflects training-set scale rather than architecture; the released corpus and benchmark make a matched-data comparison possible.

The same ranking holds on external data Aiki-HLA never trained on. The open ensemble was scored on two sources held out of its training entirely (MHC Motif Atlas and SysteMHC Atlas), with peptide–allele pairs that recur in its training through other sources removed. On this partition it reaches per-allele median AUROC **0.947** [0.936, 0.958] (Class I, 130 alleles) and **0.827** [0.811, 0.850] (Class II, 111 alleles) against proteome decoys, within the published 0.95–0.97 (Class I) and 0.82–0.88 (Class II) ranges. Strict leakage controls do not preclude conventional performance. The margin also remains after dropping Aiki-HLA’s own source-matched negatives and scoring only experimental rows (Methods).

### A calibrated presentation probability as the released model

The released model combines the allele-blind ligand-likeness score *p*_ligand_ (AUROC **0.90** Class I, **0.81** Class II against proteome decoys within distribution, **0.90**/**0.71** on a held-out set that is both later in time and sequence-disjoint; Methods) with Aiki-HLA’s allele-specific score *p*_binding_ as the geometric mean 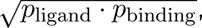, with no fitted mixing parameter. Scoring the two terms separately and multiplying yields a single model for both ligand/background screening and allele assignment (Fig. 1).

**Figure 1:**
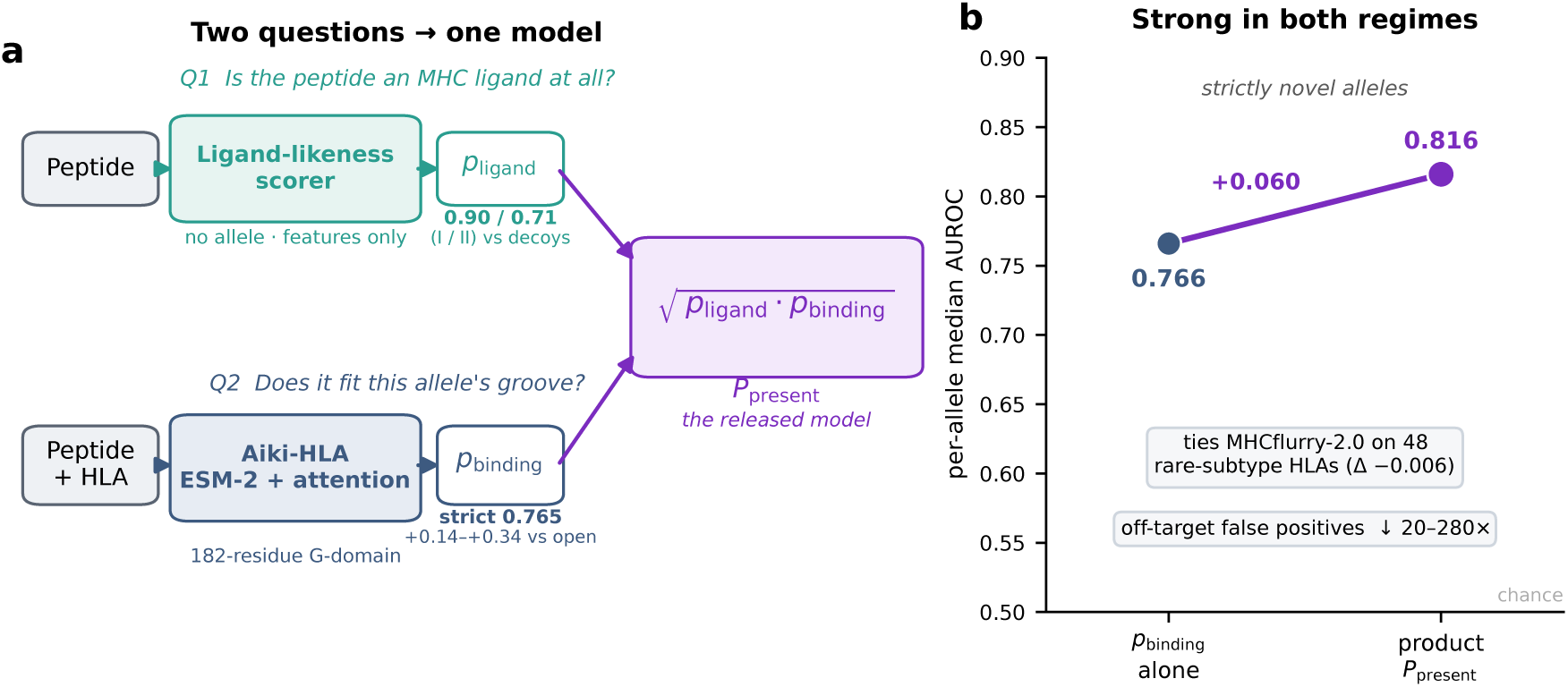
Combining a ligand-likeness score and an allele-specific score yields a presentation model for both tasks. **a**, The two components, scored separately. The ligand-likeness score *p*_ligand_ (hand-crafted peptide features, no allele input) estimates whether a peptide is an MHC ligand at all (AUROC 0.90 Class I / 0.81 Class II against proteome decoys within distribution; 0.90 / 0.71 on a held-out set that is both later in time and sequence-disjoint; Methods). Aiki-HLA’s allele-specific score *p*_binding_ (ESM-2 over the full 182-residue groove) estimates whether it fits a given allele’s groove (0.765 on novel alleles; +0.22 to +0.37 per-allele over eight comparator variants after pair-filtering each comparator’s training out of the test). The released model is their geometric mean 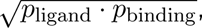, with no fitted mixing parameter. **b**, The combined score improves both tasks: it raises the novel-allele score from 0.766 to 0.816 (+0.060 over 38 comparable alleles), ties MHCflurry-2.0 across 48 rare-subtype HLAs on its own proteome-decoy task (Δ_AUROC_ = 0.006), and cuts the high-confidence off-target false-positive rate by 20–280×. (The 0.765 of panel a covers all 39 novel alleles; the 0.766 here covers the 38 with a paired combined score; Methods.)

The combined score is tested two ways. On the 39 strictly novel alleles it *raises* the lower bound from 0.766 to **0.816** (Δ = +0.060 [+0.026, +0.083]; 33 of 38 comparable alleles improve). And when screening random proteome peptides for unwanted MHC binding, it reduces the number flagged as high-confidence binders by 20–280× relative to the allele-specific score alone (below). As an external check on a proteome-decoy task that favors MHCflurry-2.0 (which was trained on proteome decoys), the composed score matches it across 48 rare-subtype Class I alleles (Δ_AUROC_ = − 0.006 [ 0.024, +0.013], 22 of 48 alleles). On this task pure *p*_binding_ alone trails MHCflurry (per-allele median 0.741 vs 0.866 on the data-rich tier; and 0.831 vs 0.904 on a separate set of 12 alleles drawn from 2024–2026 depositions that postdate MHCflurry’s training), as expected for a task MHCflurry was built for and *p*_binding_ was not. Composing *p*_ligand_ supplies that missing ligand-screening signal and brings the two level (the same composition lifts pure *p*_binding_ from 0.756 to 0.797 on the subset of these alleles with real measured non-binders). The allele-specific score *p*_binding_ is reported on its own for the strictly-novel bound (0.765) and the cross-tool margins, since each is an allele-specific measurement; the combined score is used for all other evaluations.

The factorization also matters downstream. In off-target safety screening, which scores 100,000 random human 9-mers where the hazard is a false high-confidence binder, composing with *p*_ligand_ cuts the 0.9 flag rate 20 to 280 over *p*_binding_ alone, because most random proteome peptides are non-ligands the ligand-likeness factor down-weights (Fig. S1c). The allele-specific factor serves two further tasks. It deconvolves multi-allelic mass-spectrometry panels, concentrating 46% of peptides on a single allele across 114 panels, 0.61 bits below the uniform-allele prior; the 529,129 resulting assignments feed the research-use corpus. It also scores non-classical HLA zero-shot (pooled AUROC 0.92 over 8 alleles, 0.96 once trained on them).

## Discussion

How well a pMHC predictor extrapolates to unseen MHC grooves has been hard to infer from published numbers, because the field’s data are dense on a few alleles and sparse on the rest. By holding alleles out at both the cluster and the raw-sequence level and pairing positives with provenance-matched negatives, we place that number in the high 0.7s for strictly novel alleles. That is above chance, though short of the 0.85–0.95 a conventional split returns, and the gap is a property of the split: it applies equally to our own model under a conventional split. The contribution is as much the open corpus and controls that make this measurable across both HLA classes as the predictor evaluated under them.

That predictor improves on published tools by per-allele Δ_AUROC_ = +0.22 to +0.37 (Wilcoxon *p <* 10^−9^ on every one), most on the least-studied genes, and it owes part of that margin to two design choices we describe rather than to depth of architecture: scoring both HLA classes with one model, and factoring presentation into a peptide-only ligand-likeness term and an allele-specific term whose parameter-free product is evaluated in both settings (lifting strict novel-allele AUROC from 0.766 to 0.816). The factorization is architecture-independent and applies to any predictor exposing a per-peptide score. Across both scores Aiki-HLA covers 421 (open) / 507 (research-use) HLA alleles spanning both classes (versus 171 Class I alleles in MHCflurry-2.0 and 88 Class II alleles in MixMHC2pred-v2.0, the next-broadest per-class published lists; Supplementary Table 9). The pMHC field’s foundational tools ^3,5,6,12,13,21^ and shared data infrastructure (IEDB, HLA Ligand Atlas^22^, IPD-IMGT/HLA), built over two decades, perform well where the training data are dense. The harder test for a new predictor is where data are sparse, on alleles far from any training example. That is where Aiki-HLA’s margin concentrates: it grows on the least-studied genes (HLA-C and the HLA-DQ heterodimers) and holds on a strictly novel-allele stratum where conventional benchmarks cannot measure at all. The released benchmark defines this number by its controls: novelty-aware splits at 80% peptide and 90% G-domain identity, source-matched negatives that carry no provenance signal, and a contamination audit shipped with the corpus. Applied to another tool with the same per-comparator removal of memorized training rows, it yields comparable numbers. The pair-symmetric protocol generalizes beyond pMHC: any biological prediction task whose train and test partitions share underlying entities (peptide–protein interaction, drug–target binding, antibody–epitope recognition) has the same row-level memorization confound, and the row-removal procedure is independent of the specific identity definition used.

Under these controls Aiki-HLA’s per-allele median AUROC falls in [0.71, 0.92], set by how novel a query allele is. That the same model reaches 0.947 on independent data excluded from its training (after removing pair-level overlap with its own training), within the published 0.95–0.97 range, indicates that the two measurements differ mainly in allele novelty, not in task: strict leakage controls still allow conventional-range numbers. The cross-tool margins (Δ_PA−med_ = +0.22 to +0.37, after pair-level removal of comparator-training rows) belong to the allele-assignment task; on bulk proteome screening, specialized tools perform at least as well (below).

Among the open pMHC training corpora we surveyed, this is the largest sequence-mapped set of experimental measurements assembled for training, counting only real binding-affinity and eluted-ligand observations and excluding all generated decoys: 4.66M measurements over 421 alleles in the open corpus and 5.80M over 507 alleles in the research-use corpus. For comparison, the largest single open training sets under permissive licenses are of order 10^5^–10^6^ measurements (MHCflurry 2.0 457,000; TransPHLA 502,000; NetMHCpan-4.1 ~850,000; Supplementary Table 9); the scale here comes from aggregating eleven public sources under unified allele resolution, not from any single source being larger. Large immunopeptidome resources such as the MHC Motif Atlas and SysteMHC Atlas^42^ contribute additional eluted-ligand positives to the research-use corpus; comparably-scaled primary-tissue catalogs such as PCI-DB ^41^ are possible future additions.

Our training set is 5 to 10 times larger than any comparator’s, so the cross-tool margin reflects data scale as well as architecture, and we did not run the matched-training-data ablation that would separate the two. Two observations suggest scale is a real part of it. On 48 rare-subtype HLAs scored against proteome decoys, the composed score equals MHCflurry-2.0 (Δ_AUROC_ = − 0.006, 22 of 48 alleles); and on NetMHCpan-4.2’s released held-out set, the allele-specific score ties NetMHCpan-4.1 (Δ = −0.011) on rows neither side trained on. At matched task and conventional stringency the gap narrows to parity. The released corpus and benchmark make that matched-data follow-up reproducible.

Adding more data broadens coverage but does not improve performance on alleles already covered. Extending the open corpus to the research-use corpus (~1.06 million further measurements, 86 alleles, new gene families) leaves per-allele median AUROC on the 174 shared alleles essentially unchanged ( 0.018 [ 0.026, 0.013], a third of it from a train/test decoy-method mismatch; Methods). The gain falls instead where the open subset had no training rows at all: 0.949 on 111 newly-evaluable rare subtypes, and +0.087 [+0.070, +0.100] on 234 alleles it could evaluate only at the peptide-only floor. What limits novel-allele generalization is which regions of MHC sequence space the corpus reaches, not how densely it samples the alleles it already has. Broadening coverage of HLA-C and the HLA-DQ/DPA-DPB heterodimers, not adding data on common alleles, is what would raise it.

### Limitations

Four issues bound the released evaluation. (i) All three ensembles are restricted to human MHC: pooled AUROC for the two released ensembles over 56,627 IEDB rows of rat and mouse MHC alleles is 0.558 (open) / 0.591 (research-use), so the binding rules learned here do not transfer to murine MHCs without retraining; the non-human rows present in the aggregated sources are released for a future multi-species training pass. (ii) Aiki-HLA has not been validated prospectively for neoantigen ranking; the available T-cell-response cohorts (TESLA, BNT-122) are too small (*n* = 6 and *n* = 8 patients with both classes) to assess this use. (iii) On a Class II subset of the recent (2024–2026) IEDB held-out slice the research-use ensemble underperforms the open ensemble by Δ = 0.122 [ 0.204, 0.083] (*n_a_* = 14); the cause is not decomposed between deconvolution-derived training noise, label-distribution shifts, and architecture-versus-data interaction. (iv) About 0.2% of training negatives are mis-signed (functional or stability assays that nonetheless form a complex); both ensembles score these as binders (median probability 0.36–0.73 versus a 0.10 clean-negative floor), so the noise does not affect the released metrics. A stricter automatic 90%-identity bridge purge in place of the hand-curated within-family removal lowers per-allele median AUROC by ~ 0.12 on identical evaluation rows, dominated by the removal of entire gene families under single-linkage clustering (Supplementary Table 8). The corpus, the contamination audit, the tiered-evaluation helper, and both released ensembles are available so that others can replicate on the 20 unused leave-one-cluster-out splits and re-evaluate against new IEDB submissions under the same controls.

The two design choices do different work. Factoring presentation into ligand-likeness and allele-fit gives one model for both allele assignment and proteome screening. Because Aiki-HLA’s training data are dominated by presentation events, with non-binders drawn from peptides presented by other alleles, the fitted model emphasizes allele specificity among presented ligands, whereas tools trained on proteome decoys capture the complementary ligand-versus-background signal. Multiplying the two scores recovers both, which is why we release the product rather than the allele-specific score alone.

Unifying the two HLA classes in one model gives breadth, not transfer. It scores Class I and Class II at once, a coverage the field’s class-specific tools lack. But in a controlled test the shared head shares capacity, not transferable features: adding 10% of one class to the other’s training neither helps nor consistently hurts the target class (per-allele Δ = −0.005 for Class I into Class II, +0.001 the other way; SI Table 6). The unified representation is therefore justified by simplicity and coverage, not by cross-class generalization. And on the bulk-proteome screening task the field has refined over two decades, specialized tools perform at least as well, because that task is largely ligand-likeness that simple features already capture.

## Materials and methods

### Training corpora, licensing, and ensembles

The reference predictor is released as two 3-seed dropout ensembles of identical architecture and identical training recipe, differing only in the public sources their data-extraction pipeline admits.

Throughout, allele counts refer to canonical IPD-IMGT/HLA designations (e.g. HLA-A*02:01); identical-G-domain alleles collapse to one model input, so the open corpus’s 421 canonical alleles span 332 unique G-domain inputs and the research-use corpus’s 507 span 406. We report canonical counts as the user-facing measure of coverage.

#### Open ensemble

4,662,506 experimental peptide-MHC measurements aggregated from 11 public sources whose licenses permit unrestricted redistribution, including commercial use (CC-BY-4.0, GPL-3.0, MIT, CC-BY-ND-4.0). Restricted at extraction to classical human HLA-A/B/C, HLA-DR, HLA-DP and HLA-DQ; 421 alleles after the source-matched negative-balancing step. The pre-balancing aggregated count is 4,935,914 rows over 495 alleles, of which 130,261 are curated-derived (extraction-time inferred from qualitative labels) and the balance is direct experimental measurement. 99/1 train/val partition stratified by MHC class × source; 7,078,120 training rows including source-matched cross-allele negatives. This is the cross-tool comparator subset and the unrestricted, commercial-use-permitted release.

#### Research-use ensemble

A strict superset that additionally incorporates two CC-BY-NC sources (MHC Motif Atlas, SysteMHC Atlas) contributing ~1.06M measurements, an academic-only source (MHCnuggets), 529,129 (peptide, predicted-allele) pairs from model-based deconvolution of multi-allelic mass-spectrometry data (Application demonstrations), non-classical HLA (HLA-E, F, G), and MHC class I-related molecules (MICA, MICB); these categories are deferred from the open extraction. 5,796,339 experimental measurements, 507 alleles; 98/1/1 train/val/test partition with the test partition exempting the cross-partition-overlap audit modes that the interpolation validation partition is designed to share. Released under a research-use license reflecting the CC-BY-NC and academic-only fractions.

#### Bridge-purged evaluation ensemble

A third ensemble, recipe-identical to the open ensemble but trained on a subset of the open corpus with 30 within-supertype training alleles at 0.85–0.92 raw G-domain identity to each strict-stratum test allele removed before training. This bridge purge prevents within-supertype interpolation from inflating the cluster-novel lower bound. The evaluation ensemble is not a production release. It is the model used to estimate the 0.765 lower bound. Its relation to the open and research-use ensembles is that all share the same architecture and recipe, so any model-level difference is bounded by the training rows used for each one.

### Metrics

Throughout, *per-allele median AUROC* (PA-med) denotes the median, over alleles in the named stratum, of the area under each allele’s receiver-operating- characteristic curve; *row-pooled AUROC* denotes the AUROC computed once over all rows in the stratum, ignoring allele membership. The two carry different fairness assumptions. PA-med is balanced across alleles; row-pooled weights by per-allele row count. We report both for every cross-tool comparison. *Top-100 precision* is the fraction of the 100 highest-scored test rows that are real binders. *Brier score* is the mean squared difference between predicted probability and observed label, computed on the 120K binding-affinity experimental test rows. *Expected calibration error (ECE)* is the probability-weighted mean absolute calibration gap, binned in 15 equal-mass deciles^23^. *Bootstrap 95% confidence intervals* are computed by resampling alleles within each stratum 1,000 times ^24^; the same resampling supports the Wilcoxon signed-rank test for the per-allele paired-difference summaries. Reported *p*-values are nominal and are not corrected for multiple comparisons across strata, comparators, or alleles; the per-test hypotheses (each row of Table 2 and each strict-stratum source partition) are pre-specified by the leakage-controlled paired-row protocol and the four named evaluation buckets rather than discovered post-hoc.

### Data sources and leakage controls

We aggregated ~60M pMHC observations from 11 public sources: IEDB^22^, HLA Ligand Atlas, CEDAR, NetMHCpan training data, NetMHCIIpan training data, NetMHCstabpan training data, TransPHLA positives, and VDJdb under commercial-compatible licenses; plus MHC Motif Atlas, SysteMHC Atlas, and MHCnuggets, excluded at extraction under non-commercial / academic-only licenses. A unified allele-to-sequence database (37,480 entries from IPD-IMGT/HLA, IPD-MHC, UniProt, NCBI) implements direct IMGT matching, compound Class II heterodimer chain splitting, and serotype recovery, recovering 2.46M Class II compound-heterodimer rows that direct matching alone would discard. CEDAR rows are ~100% IEDB-contained at the peptide × allele level and are deduplicated into IEDB during corpus construction. Of the 60M raw aggregated rows, 4.94M experimental measurements survive the filters and allele-to-sequence resolution for the commercial-use (open) corpus; adding the non-commercial and deconvolution-derived sources brings the research-use corpus to 5.80M (Results). The rest, dominated by third-party generated decoys excluded at extraction, are released as named hold-out artifacts. Per-source row counts, licensing, and the full funnel are in Supplementary Table 3; a non-MHC shortcut diagnostic confirms that every source × class cell carries label signal from provenance alone (*p <* 10^−3^), motivating the negative-generation validator below.

At training entry an automated audit re-verifies the corpus against the contamination modes the benchmark controls (chiefly a decoy mislabeled as a measured positive, or a peptide–allele pair appearing as both) and halts the run if any is found. The audit is included with the corpus so that a rebuilt or extended corpus can be re-certified to the same standard; the released and evaluation corpora pass it.

### Negative generation

Many pMHC sources are positive-only (IEDB mass-spectrometry, TransPHLA positives, HLA Ligand Atlas ligandomes), so a classifier on the raw aggregation can score near-perfectly by predicting a source’s predominant label rather than binding. We generate negatives by one source-matched procedure with three layers, designed to make source, length, and provenance non-predictive of the label.

*(i) Source-matched cross-allele negatives* (the bulk of generated negatives). For each (source, allele, peptide length) group, candidates are drawn from a different MHC G-domain cluster than the source allele, length-matched, within the same MHC class, with the (allele, peptide) pair never appearing as a positive at that allele anywhere in the corpus. Each negative inherits the source positive’s source, measurement-type and provenance fields, making those features non-predictive at the row level. Hard-negative weighting gives higher weight to peptides observed as positives in many other MHC clusters.
*(ii) Cross-class deficit fallback.* When within-class candidates are exhausted at a length (typical for HLA-C and HLA-DQ heterodimers), we fall back to cross-class peptides (for example a Class I 9-mer as a negative for a Class II allele). These fill per-allele ratio gaps without synthesizing new peptides, which would create class-pure decoys and break the source-matching.
*(iii) Pos-only memorization guard.* For peptides that are positive somewhere in a split but never negative anywhere in it, one distant-cluster cross-allele negative is added, so no peptide is memorizable by its label alone.

The generator also implements anchor-preserving shuffling and proteome-fragment negatives, but neither is used in the released corpora: in development both introduced an exploitable composition signature (a peptide-only classifier separated their decoys from real negatives), so we retained only the source-matched procedure above.

A built-in validator gates the build: a metadata-only logistic regression (length, source, measurement type) must score AUROC *<* 0.60 per feature and *<* 0.65 combined, or the build fails. A peptide-only ESM-2 probe quantifies the residual origin signature (AUROC 0.67 corpus-wide, 0.80 on Class II×IEDB). The strict novel-allele stratum excludes generated negatives entirely (experimental rows only), isolating the binding signal from any decoy signature; as a companion check, the open ensemble itself separates experimental from generated negatives only at chance (per-allele median 0.516 over 26 alleles, none above 0.70; Supplementary Table 7), and the full-vs-experimental-only pooled Δ is 0.031, indicating that decoys slightly lower the headline rather than inflate it. On three nested decoy-exclusion slices (full / EL-removed / BA-only) the evaluation ensemble achieves PA-med 0.763 / 0.832 / 0.715 on the broad 273-allele stratum; the strict 39-allele stratum is experimental-only (no generated decoys) by construction.

### Cluster-aware splitting and strict novel-allele stratum

The field’s standard novel-allele splits typically control for the allele axis at a 95%-identity threshold that does not separate cluster-novel from cluster-mate alleles^18,29^ (73–83% of typical test alleles fall above 0.95 G-domain identity to a training allele in our audit). We split on *both* peptide and MHC axes simultaneously: MHC G-domain sequences are clustered at 90% identity via MMseqs2^10^ (single-linkage), and bridge alleles (those that link otherwise-separate clusters at the 0.85–0.90 identity band) are purged from training to break the linkage and enforce stricter cluster separation in test. Peptides are clustered independently at 80% identity. The 3×3 grid generates four evaluation buckets: trivial, novel-peptide, novel-allele, and both-novel. A second, tighter 0.95 within-cluster identity threshold is used only by the strict-stratum drop-set selection (below) to identify the cluster mates of a named held allele; this 0.95 threshold is *not* the cluster threshold and does not appear in the 90%-clustered split structure.

#### Cluster-level versus sequence-identity disjointness

Single-linkage 90%-identity clustering separates test alleles from their nearest training neighbor at the cluster-label level but not always at the raw-sequence level: a representative-greedy heuristic can place two near-identical alleles in different clusters. An independent re-clustering audit finds 153 of 372 broad-stratum test alleles have a training allele at ≥0.90 raw G-domain identity, 56 at ≥0.99 (e.g. HLA-A*02:01 test vs HLA-A*02:12 train, 0.994). We disclose this and address it two ways. (a) The headline lower bound uses only the sub-stratum below 0.90 raw identity (39 alleles, 140,073 experimental rows, PA-med 0.765 [0.691, 0.797]), which is cluster-and sequence-disjoint. (b) The 0.90–0.93 band above it is reported separately as a near-twin stress test (20 alleles, 50,371 rows, PA-med 0.661). The audit code, and leave-one-class-out partitions for the strictest disjointness (max test-to-train identity ≤0.78 by construction), are released (Data availability).

#### Bridge-purged training corpus

The evaluation ensemble’s training corpus (the cluster-novel lower-bound estimator) purges 30 within-supertype bridge alleles whose raw G-domain identity to a held strict-stratum test allele falls in the 0.85–0.92 band; leaving these in training would let the model interpolate between near-identical neighbors instead of extrapolating. The purge fixes at build time, applies to train and val, and leaves test unchanged. It removes 3.4% of training rows (4.94M → 4.77M), and every held test allele’s nearest training neighbor is then at *<*0.93 G-domain identity by construction. The 39-allele headline sub-stratum (Table 1) is selected post-hoc against this bridge-purged set at the stricter *<*0.90 threshold so the stratum is sequence-disjoint, not merely cluster-label-disjoint. The two production ensembles (open and research-use) train on the respective full non-purged corpora.

### Model, training, and ensemble

The architecture feeds unmodified ESM-2 650M^4,16,26^ weights (esm2_t33_650M_UR50D) embeddings (dim 1280, cached) of two inputs: the peptide (length 8–25) and the MHC G-domain as one 182-residue sequence: Class I as the natural *α*1 + *α*2 groove (90 + 92), Class II as the 172–181-residue *α*1 + *β*1 groove padded to 182 with a short tail from the same *β* chain (just beyond the groove). The shared 182-residue shape lets one head score both classes; a mask token fills missing residues in 0.04% of Class II and 0.13% of Class I rows. Using the full G-domain rather than the 34-residue pseudosequence^27,28^ is the substantive departure from prior Class II predictors^7–9,13,30–32;^ recent both-class tools instead use pseudosequence + class-specific heads^38^, a single-chain interaction grammar ^37^, or pMHC–TCR unification ^39^.

Embeddings are projected to *d*_model_ = 256 and passed through one pre-norm self-attention layer (8 heads, ~3.9M trainable params), position-biased peptide pooling, multi-head MHC pooling, and a 3-layer MLP head (dropout 0.2) with two weight-shared branches: a binary classifier (the released binder probability) and a regression head on log IC_50_ for the 18.7% of rows with a continuous affinity measurement. Training: focal loss (*γ* = 2.0, *α* = 0.43) on binary rows; an *inequality MSE* on the regression branch penalizing only predictions beyond the assay’s reported bound (e.g. *>* 50,000 nM treated as a lower bound); label smoothing 0.05; AdamW (*η* = 10^−4^, weight decay 10^−2^); AMP FP16; batch 256; 6 epochs; patience 2; gradient clipping *g* _2_ 1; 5% linear warmup then cosine decay to 10% of peak; mask-aware pooling; three seeds averaged at inference.

#### Production ensembles and lower-bound estimator

Each production ensemble (*open*, *research-use*) trains on 99% of its corpus (bridges included), 1% held out for monitoring; early stopping never triggered (val AUROC rose monotonically through the locked 6 epochs). Because both production ensembles have seen the strict stratum during training, all lower-bound numbers come instead from the recipe-identical *evaluation ensemble* on the bridge-purged corpus (test held out), which lower-bounds either production ensemble on strictly novel queries (the standard held-out-fold-estimator vs trained-on-everything-production relation). Upper-bound numbers come from a production ensemble’s own 1% held-out partition; cross-tool numbers from the open ensemble’s 1% partition. A frozen-ESM head with 0.2 dropout was selected from a 9-axis sweep (depth, attention variants, sampling, regularization, loss calibration, negative resampling, and four representation-flexibility paths: LoRA at two ranks, full ESM-2 fine-tuning, adapter-warmstart); no trainable-embedding variant exceeded it by a defensible margin (Supplementary Fig. S4). On the 174 alleles both production ensembles evaluate, the open-vs-research-use delta is 0.018 [ 0.026, 0.013] ( 0.011 residual after the experimental-only control), attributing about a third to a train/test decoy-method mismatch affecting ~0.1% of research-use training. The effect is bounded by the ~0.007 open/research-use residual and does not enter the strict-stratum, upper-bound, or composite headlines.

### Architecture ablation and trivial-floor baselines

The architecture ladder on the both-novel Class I bucket (70 alleles; *k*-NN 0.512 → pan-LR 0.526 → pan-MLP 0.631 → attention-head ensemble 0.668) is in Supplementary Fig. S4. On the strict 39-allele stratum the strongest peptide-only baseline reaches 0.493 (length-only 0.509), so the 0.272 lift to 0.765 is MHC-context-driven. A same-allele metadata probe (length+source+assay) ^11,19^ reaches 0.619 corpus-wide on the broad stratum, confirming that heterogeneous source negatives carry source-correlated structure; this effect is absent by construction on the strict stratum, which uses experimental negatives only and enforces peptide-cluster novelty.

### Ligand-likeness scorer

Two class-specific peptide-only *ligand-likeness scorers* (Class I and Class II) score *p*_ligand_, the probability that a peptide is MHC-shaped at all, independent of any specific allele. Features are hand-crafted: amino-acid composition, peptide length, hydropathy mean and standard deviation, net and absolute charge, fractional hydrophobic / acidic / basic content, and N- and C-terminal three-residue one-hot encodings (anchor-capturing, length-agnostic). No learned embedding and no allele input are used. Each scorer is a logistic regression stacked with a HistGradientBoosting classifier, fit on positive eluted ligands versus a blend of length-matched random human-proteome fragments and anchor-shuffled positives (the latter to emphasize positional anchors rather than bulk composition); recent-IEDB test peptides are excluded from ligand-likeness training. Held-out AUROC (random 80/20 split of the training pool): Class I 0.869 (0.900 against pure proteome decoys), Class II 0.718 (0.813 against pure proteome decoys). On a stricter held-out that is both temporal (recent-IEDB positives postdating the training pool) and peptide-cluster-disjoint (MMseqs2 80% identity), the Class I scorer is essentially unchanged (0.902 against proteome decoys) while the Class II scorer falls to 0.708, so the same in-distribution inflation this manuscript documents for allele-specific prediction also affects the peptide-only factor. A data-scaling analysis over the full 1.70M-peptide ligand union and an ESM-2 mean-pool capacity probe confirm this Class II ceiling is not relieved by more positive data (the scorer saturates by 60,000 pos-itives per class) or by learned embeddings (both reach ≈0.70 against proteome decoys), which is why the released product relies on the allele-specific factor *p*_binding_ for Class II. The composed presentation probability is the equal-weight geometric mean of the ligand-likeness and allele-specific binding scores, 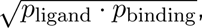, with no parameter fit on evaluation data. Both ligand-likeness scorers and the composition rule are released alongside the Aiki-HLA ensembles (Code availability); the composed product 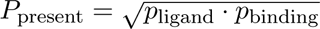 is the primary released model, with the allele-specific score *p*_binding_ exposed as a component for allele-specific evaluation (cross-tool comparison, cluster-novel discrimination).

### Cross-tool comparison protocol

Comparisons are computed on the open ensemble’s 1% held-out validation partition (70,781 rows, stratified by MHC class × source; held out from optimization and from early-stopping decisions, which did not trigger). The partition is in-distribution for both Aiki-HLA and each comparator: it is randomly sampled from the full open corpus, and both Aiki-HLA and the comparators draw on overlapping IEDB-derived data. For each comparator we use the published model weights without retraining ^5,6,12,13^ and restrict to rows whose allele and peptide length the comparator supports; pan-species predictors^3,28^ are scored on their human-HLA subset (no species or licensing confound). The research-use ensemble trains on non-classical HLA the comparators do not support; using it here would require a comparator-side allele filter we avoid.

#### Pair-level leakage filter (symmetric)

The 1% partition is row-level held out from Aiki-HLA’s training but is in-distribution for the comparators, and on the comparator side many test rows are literal training rows: without filtering, up to a third of the test would favor comparator memorization. For every comparator we therefore remove from the test subset every (peptide, allele) pair that appears in that comparator’s training data. The training pair sets used:

- MHCflurry 2.0: full curated training (curated_training_data.csv.bz2 from the MHCflurry distribution), ∼949,129 unique pairs.
- TransPHLA: 5-fold released train_data_fold[0-4].csv + val_data_fold[0-4].csv union, ∼1.86M pairs.
- NetMHCpan-4.1 EL: shipped c00[0-4]_el CV partitions (DTU NAR supplementary). NetMHC-pan ships these with a sample-id column that, for multi-allelic mass-spec entries, refers to a sample (e.g. Apher6 = HLA-A*03:01 / A*29:02 / B*44:02 / B*44:03 / C*12:03 / C*16:01) rather than a single allele; we expand each such row to its constituent alleles via the shipped allelelist file, yielding ∼45.4M unique (peptide, allele) pairs.
- NetMHCpan-4.1 BA: shipped c00[0-4]_ba partitions, 208,093 pairs (alleles named directly per row).
- NetMHCIIpan-4.1: NetMHCIIpan-4.0’s train_EL[1-5] and train_BA[1-5] (DTU NAR sup-plementary; same training distribution inherited by 4.1 and a near-superset of any subsequent 4.3 update), with EL sample-id expansion via the shipped allelelist; ∼24.7M unique pairs total.
- BigMHC EL: own training not shipped with the package (Mendeley deposit). We use MHCflurry-2.0’s curated training as a conservative under-removing proxy since BigMHC trains on a superset (MHCflurry curated + extra MS); footnoted (‡).
- MixMHC2pred v2.0: own training peptide list not shipped with the binary distribution (it ships PWMs only; raw training data is in the Racle et al. 2023 Immunity supplementary Mendeley deposit). We apply NetMHCIIpan-4.0’s pair set as a conservative proxy because the two predictors draw on the same public Class II MS + IEDB peptide pool (HLA Ligand Atlas, IEDB MS, lab MS releases); under- removing by exactly the MixMHC2pred-only training rows biases the resulting Δ downward (against Aiki-HLA); footnoted (§).

The same pair-level filter applied symmetrically against Aiki-HLA’s own training would remove no rows from this partition because the 1% val is itself the held-out 1% of Aiki-HLA’s training corpus (row-level disjoint by construction); the filter is therefore a one-sided correction restoring symmetry to a comparison that the unfiltered protocol biased against Aiki-HLA. Peptide-only filters (removing rows where the peptide appeared in the comparator’s training data with any allele) over-correct in the opposite direction and are not used.

#### Decoy-advantage control

Dropping all generated-decoy rows from the unfiltered 1% val partition (the no-pair-filter scaffold) controls for whether Aiki-HLA’s training on source-matched negatives gives it an unfair advantage: on that scaffold the MixMHC2pred v2.0 delta is essentially unchanged (+0.303 vs +0.340 unfiltered); the MHCflurry and BigMHC EL deltas drop ~ 4 pp. Under the pair-filter regime (and so removing Aiki-HLA’s generated negatives simultaneously) the Class I deltas are +0.306 [+0.189, +0.450] (MHCflurry), +0.255 [+0.202, +0.348] (TransPHLA), +0.254 [+0.186, +0.389] (BigMHC EL) on the surviving experimental-only subset: the pair-filtered Table 2 results stay within bootstrap CI after generated-negative removal.

#### Held-out neither-side-trained-on control

As an external fairness reference, both Aiki-HLA and NetMHCpan-4.1 EL (via IEDB API) are scored on the NetMHCpan-4.2-released eval pool (iedb_test + cedar_test, 12,107 Class I rows; held out from NetMHCpan-4.1 training by the 4.2 authors, and held out from Aiki-HLA’s training after an additional (peptide, allele) pair-filter against Aiki-HLA’s train+val pair set, leaving 2,019 paired rows over 13 evaluable alleles). On this strictly held-out subset Aiki-HLA (open-ensemble) and NetMHCpan-4.1 EL are statistically tied: Δ_PA−med_ = − 0.011 [ 0.086, +0.028], 6 of 13 alleles favor Aiki-HLA, both at per-allele median AUROC ~ 0.64. The Class II analogue is under-powered (NetMHCIIpan-4.3’s eval leaves a median of 3 positives per allele after the pair-filter).

## Acknowledgments

We thank the maintainers of IEDB, HLA Ligand Atlas, and CEDAR for making high-quality immunological data freely available, and those of IPD-IMGT/HLA and UniProt for the reference allele and protein sequences, and the developers of MHCflurry, BigMHC, TransPHLA-AOMP, MixMHC2pred, NetMHC-pan, and NetMHCIIpan, whose released predictors and training data made the cross-tool benchmark possible. We thank Prof. Lawrence Stern, Dr. Shankar Shastry, Dr. Eswar Iyer, Marta Chronowska and Radheesh Sharma Meda for helpful discussions.

## Use of AI tools

Anthropic Claude and OpenAI ChatGPT were used for code assistance and manuscript editing. All pipelines were specified, run, and validated by the author, and every reported number comes from deterministic, version-controlled scripts released with the manuscript (Code availability). The manuscript text was drafted and directed by the author. No quantitative result, figure, table value, citation, or scientific claim was generated by an AI tool. The study design, analyses, interpretation, and conclusions are the author’s, and each numerical claim was checked against its source artifact. No AI tool is an author or meets authorship criteria.

## Author contributions

V.M. conceived the study, designed and implemented the predictors and the evaluation framework, performed the experiments and statistical analyses, prepared the figures, and wrote the manuscript.

## Data availability

A Zenodo deposit (concept DOI 10.5281/zenodo.20520819; this v1.0.0 release 10.5281/zen-odo.20520820) contains the following artifacts under CC-BY-4.0. (1) Two training-corpus parquet files restricted to the redistributable licenses (IEDB and HLA Ligand Atlas): corpus_experimental.parquet (the experimental subset of the released 4.94M-row corpus) and corpus_balanced.parquet (the same subset with generated negatives). (2) For the remaining license-restricted sources (NetMHCpan, NetMHCIIpan, NetMHCstabpan, TransPHLA, VDJdb, BigMHC, MHCflurry, plus parsed-and-deduplicated CEDAR, and the three non-commercial sources excluded from the open corpus but included in the research-use corpus: MHC Motif Atlas, SysteMHC Atlas, MHCnuggets), per-source manifests are provided (peptide_hash, allele, measurement_type, source_record_id, fetch_script) so readers can re-pull from upstream and rebuild the rows under each source’s own license terms. (3) Train, validation, and test splits, the 39-allele cluster- and sequence-disjoint strict stratum (membership list and per-allele row counts), and the four bucket assignments (trivial, novel-peptide, novel-allele, both-novel). (4) 20 leave-one-cluster-out splits released to enable prospective multi-cluster replication. (5) Two 3-seed ensemble checkpoint sets, SHA-256-pinned in the deposit’s checkpoints/MANIFEST.json: the *open* ensemble (CC-BY-4.0; corresponds to the fully-redistributable 4.94M-row corpus in (1)) and the *research-use* ensemble (CC-BY-4.0 model weights, research-use license reflecting the CC-BY-NC and academic-only fractions of its training corpus). The research-use-ensemble training corpus is released as a manifest-only artifact for the license-restricted fraction; a rebuild_research_use.py script re-derives the research-use corpus from upstream under each source’s own license. Users requiring a stricter interpretation of redistribution restrictions can re-derive the research-use ensemble locally from that script. (6) The MHC G-domain ESM-2 embedding binary (357 MB) and embedding indices, plus a regeneration script for the 101 GB peptide embedding store which exceeds Zenodo’s deposit size limit. (7) Cross-tool paired-prediction CSVs for the head-to-head subsets (Aiki-HLA vs MHCflurry 2.0, TransPHLA, BigMHC EL, MixMHC2pred v2.0). (8) Application-benchmark inputs (TESLA cohort ^14^, BioNTech BNT-122 candidates, SKEMPI v2^15^). (9) **Multi-allelic deconvolution release:** 529,129 unique (peptide, predicted-allele) pairs from model-based deconvolution of 2.04 million experimental multi-allelic mass-spectrometry observations using the 3-seed open ensemble, parquet-deposited as deconvolution_v38/; per-row confidence band (single-candidate / low / mid / high) and runner-up allele included to support downstream filtering. The deduplication ratio (3.85×) follows from the same peptide being observed in multiple cell lines and assigned to the same dominant allele in each. The upstream multi-allelic tag in our aggregated corpus covers a larger 30 M-row pool (Methods, “Negative generation”), but 28 M of those are generated decoys that inherit the tag through provenance; only 2.05 M are experimental positives. (10) Result JSONs backing every numerical claim in this manuscript and an executable verification script (verify_paper_numbers.py). (11) Per-split independence-audit JSONs for every released split design (default, bridge-purged sister, leave-one-cluster-out, leave-one-class-out, 5-fold CV), produced by re-clustering each split’s train+test allele set from raw sequences without inheriting our cluster IDs; each audit records the per-test-allele max identity to its nearest training allele plus a histogram and a worst-offender list. Together with (10), these support an external reader applying either cluster-level or sequence-identity disjointness to any prediction file deposited with this work. DATA_SOURCES.md within the deposit documents the per-source provenance and license decisions. Public upstream source data: IEDB (https://www.iedb.org) ^22^, HLA Ligand At-las (https://hla-ligand-atlas.org), CEDAR (https://cedar.iedb.org), NetMHCpan training data (https://services.healthtech.dtu.dk/services/NetMHCpan-4.1/), NetMHCIIpan train-ing data (https://services.healthtech.dtu.dk/services/NetMHCIIpan-4.0/), NetMHCstab-pan training data, TransPHLA positives^6^, VDJdb (https://vdjdb.cdr3.net). Comparator soft-ware used for cross-tool evaluation but not folded into our corpus: MHCflurry 2.0^12^, BigMHC EL^5^, TransPHLA-AOMP, and MixMHC2pred v2.0^13^. Allele sequences from IPD-IMGT/HLA (https://www.ebi.ac.uk/ipd/imgt/hla/), IPD-MHC (https://www.ebi.ac.uk/ipd/mhc/), UniProt (https://www.uniprot.org), and NCBI (https://www.ncbi.nlm.nih.gov). SKEMPI v2 source data at https://life.bsc.es/pid/skempi2.

## Code availability

Source code is released on GitHub at https://github.com/aikium-public/aiki-hla under the Apache-2.0 license; the permanent Zenodo snapshot DOI of the GitHub release is 10.5281/zen-odo.20520820 (concept DOI 10.5281/zenodo.20520819 for the latest version). Both the open and research-use 3-seed ensemble checkpoints are released under CC-BY-4.0 (model weights) alongside the corpus on Zenodo; the research-use-ensemble *training corpus* carries a research-use license inherited from its CC-BY-NC and academic-only fractions, and is shipped as a per-source manifest with a re-build script (Data availability). The Python package provides inference, scoring, and tiered evaluation (pip install aiki-hla; command-line entry aiki-hla score –peptides X –alleles Y). A re-producible cloud-inference pipeline (Modal apps) covers the leakage-controlled corpus build, the runtime audit, and full training-recipe reproduction. The figure-generation scripts and the executable verification script (validation/reproduce_paper_numbers.py, which re-derives every headline numerical claim from the deposited artifacts within documented tolerance and validates input-file SHA-256 hashes) are included in the code repository.

## Supplementary Figures

**Figure S1:**
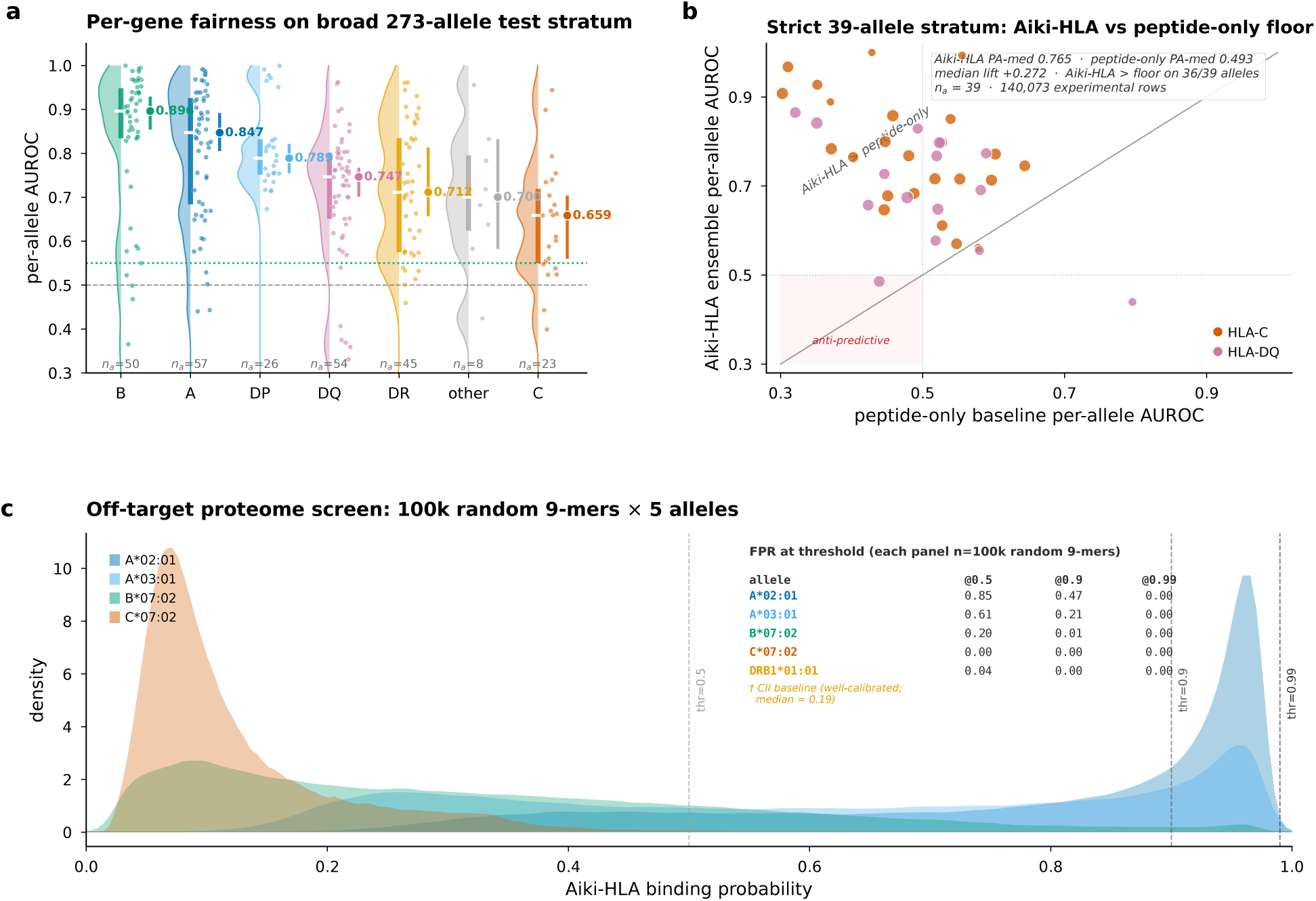
Aiki-HLA across HLA genes, against a peptide-only baseline, and in off-target screening. **a**, Performance is consistent across HLA genes on the broad 273-allele test set. HLA-B is strongest (median AUROC 0.896); HLA-C is the weakest (0.659). **b**, On the 39 strictly novel alleles, Aiki-HLA (vertical axis) outperforms a peptide-only baseline (horizontal axis) on 36 of 39, with a per-allele median gain of +0.272; the binding signal comes from the MHC, not the peptide alone. Marker size scales with test-row count, color with HLA gene. **c**, Off-target screen of 100,000 random human proteome 9-mers against four Class I alleles and HLA-DRB1*01:01: fewer than 0.001 are flagged at high confidence (≥ 0.99), the setting relevant to TCR-mimetic safety. Background levels differ across alleles (HLA-DRB1*01:01 median 0.19 vs HLA-A*02:01 0.89); composing with the ligand-likeness score further cuts the ≥ 0.9 flag rate 20 to 280× (Methods).

**Figure S2:**
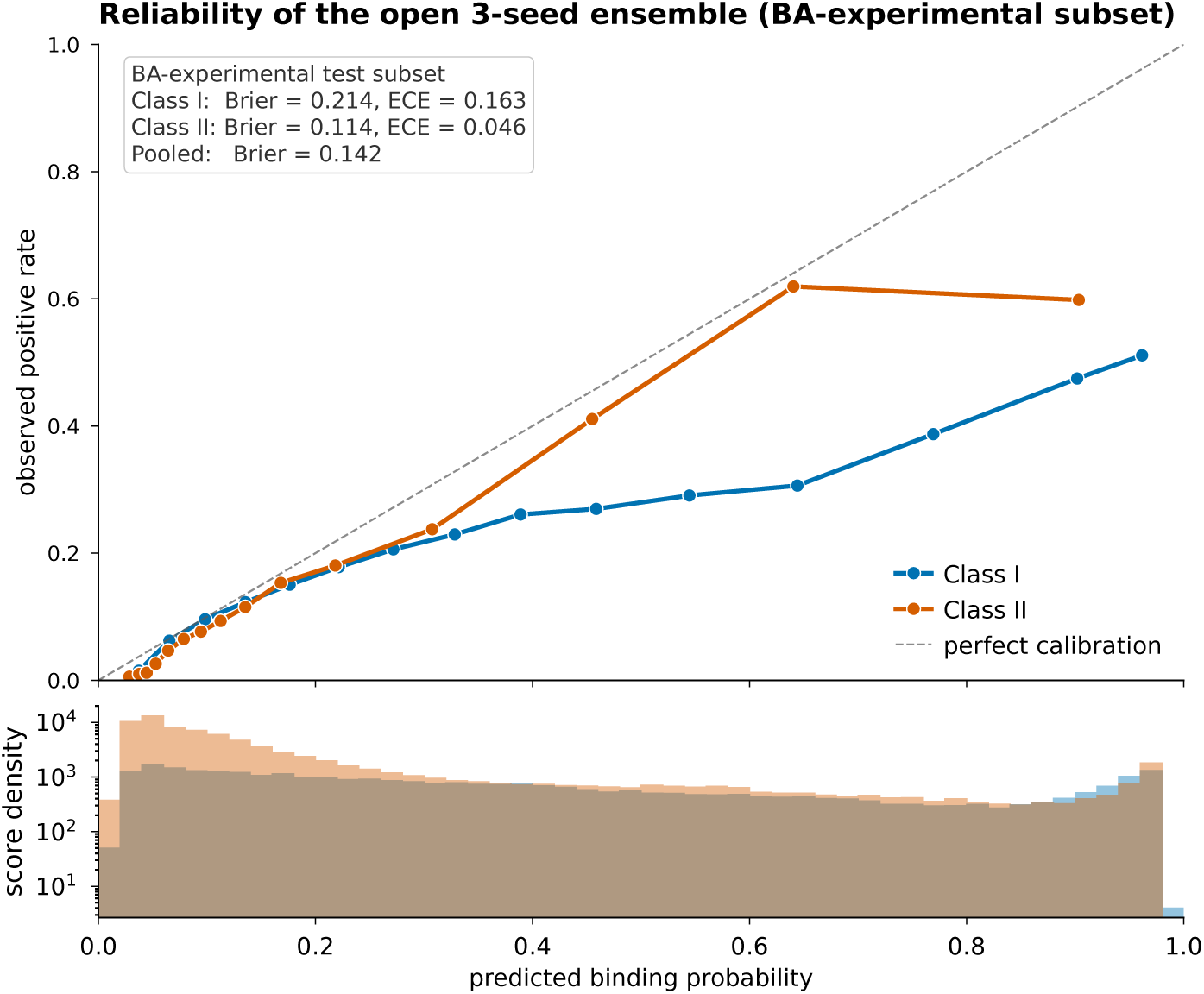
Open-ensemble probabilities are well calibrated for Class II but overconfident for Class. **I.** Predicted probability versus observed binding rate (15 equal-size bins), separately for Class I and Class II, on the binding-affinity test subset (*n* = 120,342); the diagonal is perfect calibration and the bottom strip shows score density. The Class I curve falls below the diagonal above *p* = 0.5 (overconfident), while Class II tracks the diagonal closely. Per-class Brier and calibration-error values are in the main text. For probability-thresholded use, rescale on the target allele set or set per-allele thresholds.

**Figure S3:**
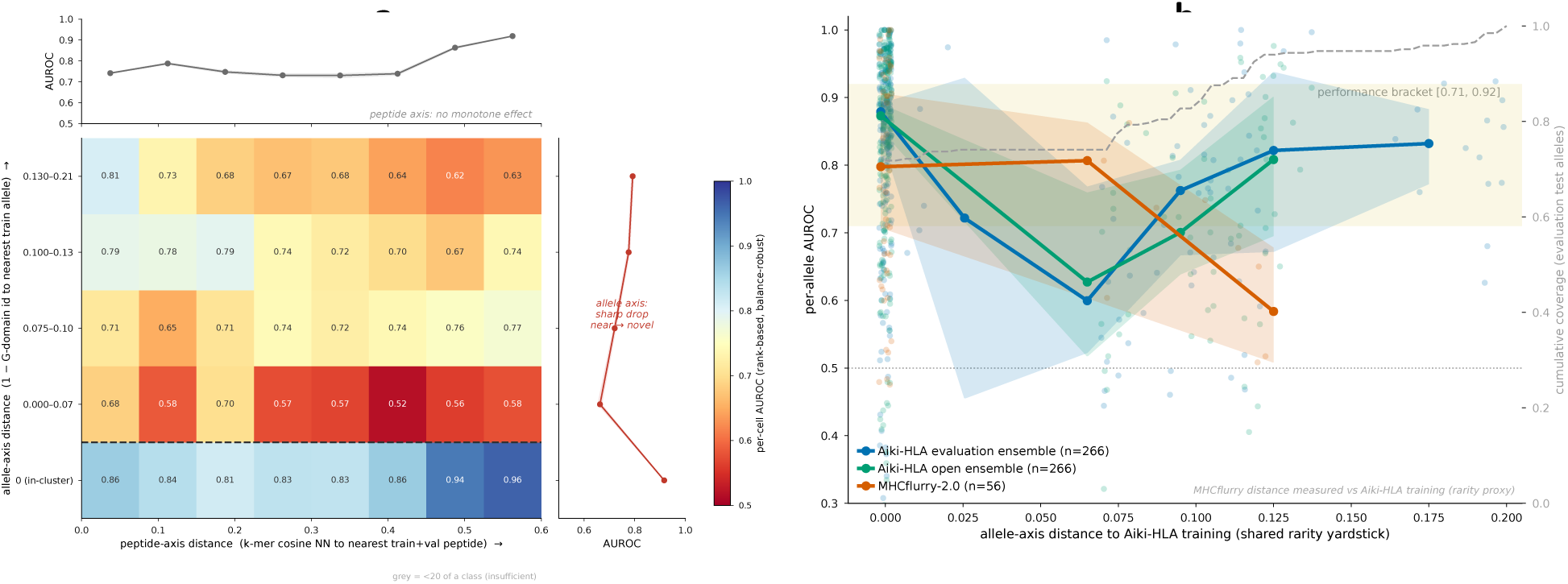
Allele distance, not peptide distance, explains most of the generalization drop, and the same pattern limits every tool. **a**, AUROC of the Aiki-HLA evaluation ensemble (*n* = 1,372,156 experimental rows) for test rows grouped by their distance to the nearest training peptide (horizontal) and nearest training allele (vertical; 1 G-domain identity). Accuracy changes sharply with allele distance (alleles close to training, bottom row, reach 0.81–0.96 regardless of peptide distance, while any novel-allele row drops to 0.52–0.81) and barely varies with peptide distance. The 727,000 training peptides already cover peptide space densely, so the MHC groove is the main axis on which generalization degrades; the hardest band is HLA-C. Marginals give AUROC versus each axis (bootstrap 95% CI); gray cells have fewer than 20 of a class. Peptide distance uses a 3-mer sequence metric because mean-pooled embeddings of short peptides have too little dynamic range. **b**, Per-allele AUROC versus distance to the nearest Aiki-HLA training allele, for the Aiki-HLA evaluation and open ensembles and MHCflurry-2.0; all three decline together as allele distance grows, indicating that the limit is set by the data rather than by any one model. Lines are per-bin medians with 95% bootstrap CI ribbons; the yellow band marks the [0.71, 0.92] bracket; the dashed line (right axis) is the cumulative fraction of test alleles within distance *x*. Allele distance is measured against Aiki-HLA’s training set as a common rarity scale for all tools.

**Figure S4:**
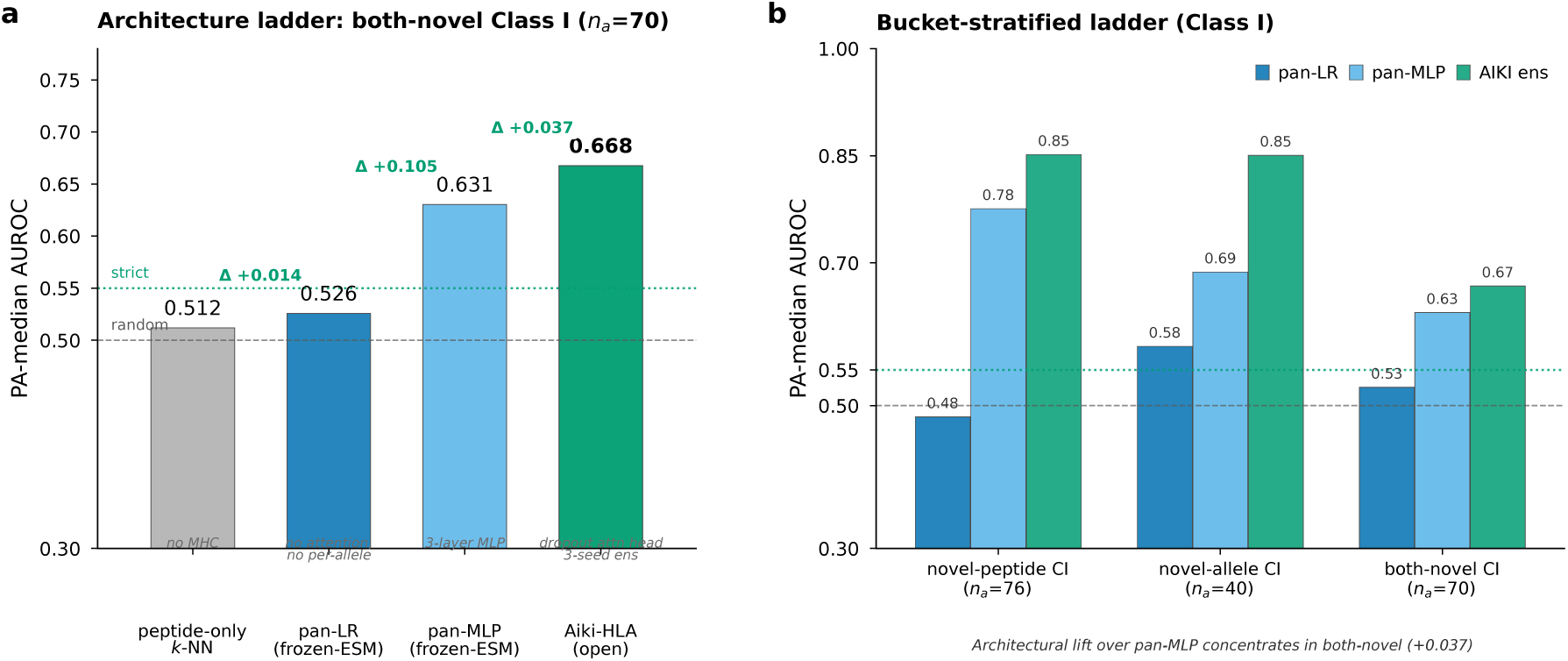
Reading the MHC groove with an attention head improves performance on novel alleles. **a**, On the hardest (both-novel) Class I alleles (*n_a_* = 70), accuracy climbs as the model gains MHC context: a peptide-only baseline (no MHC) reaches median AUROC **0.512**, a linear model on pooled MHC features **0.526**, a 3-layer network **0.631**, and the Aiki-HLA attention-head ensemble **0.668**. Green dotted line, 0.55 reporting threshold; gray dashed, chance. **b**, Across the three novelty buckets, the attention head’s gain over the plain network is largest on the both-novel bucket (Δ = +0.037), where the simpler models fall to near chance.

## Supplementary Tables

**Table 3:**
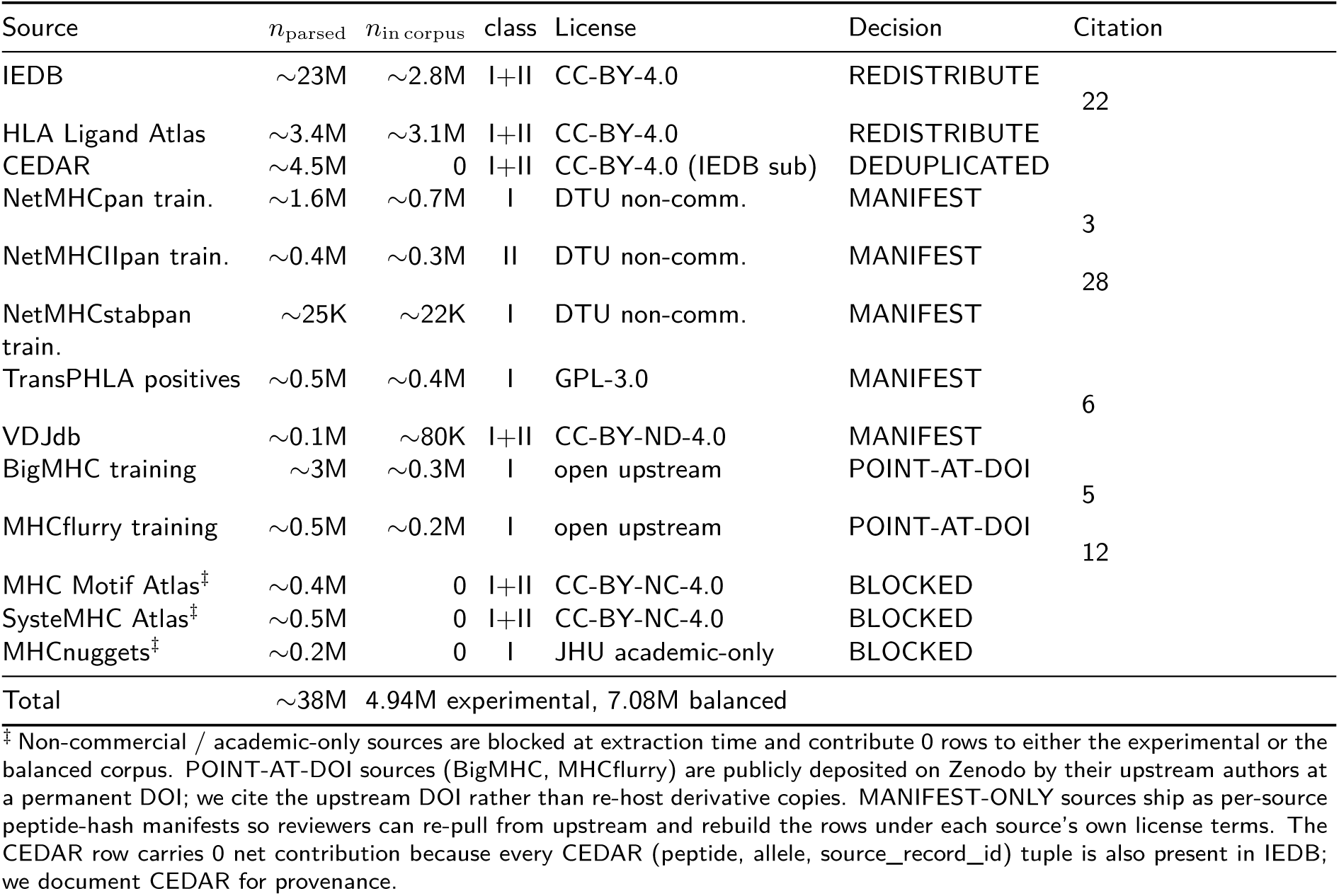
Per-source data inventory and Zenodo redistribution decisions. The aggregation parses 11 public pMHC sources; non-commercial sources are blocked at extraction (MHC Motif Atlas CC-BY-NC-4.0; SysteMHC Atlas CC-BY-NC-4.0; MHCnuggets JHU academic-only). *n*_parsed_ is rows after format harmonisation; *n*_in corpus_ is rows surviving deduplication into the final 4.94M experimental set (CEDAR is parsed but ~100% IEDB-contained, so 0 unique CEDAR rows survive). Redistribution decisions follow the corpus release plan: REDISTRIBUTE for unambiguously permissive sources (IEDB, HLA Lig-and Atlas); MANIFEST-ONLY for license-restricted or copyleft sources, in which case a per-source CSV of (peptide_hash, allele, measurement_type, source_record_id, fetch_script) is shipped so reviewers can re-pull from upstream and rebuild the corpus row-set.

**Table 4:** Per-allele cluster-distance map for 84 both-cluster-novel test alleles. For every test allele in the *both-cluster-novel* bucket of Table 1, the raw G-domain identity (position-aligned percent identity over the 182-residue platform sequence) to the nearest training allele after the bridge purge is recorded and bucketed. Confirms no allele leaks into the bridge zone after the corpus build. The bands above are the cluster-distance map; the headline PA-med 0.765 scored stratum is a *different* cut: the ≥5-positive/≥5-negative-qualifying subset of *all* 55 alleles at *<*0.90 identity (the highly_novel strict_stratum bands), i.e. 23 HLA-C + 16 HLA-DQ, not the 34 HLA-C + 5 HLA-DQ of the strict band alone. Under an alignment-based (MMseqs2) identity the HLA-DQ group stays cluster-novel (*<*0.90, median 0.89; the position-aligned 0.802–0.835 understates heterodimer identity because the variable-length *α*+*β* chains misalign position-wise), while 7 bound-ary HLA-C alleles (~0.896 position-aligned) reclassify just above 0.90; the HLA-DQ generalization claim is therefore insensitive to this metric choice, while the HLA-C boundary is metric-sensitive. Full 84-row TSV deposited on Zenodo (outputs/v37_cluster_distance/cluster_distance_map.tsv; columns allele, mhc_class, max_id_to_train, nearest_train_allele, bucket).

| Bucket | $n_{\text{alleles}}$ | Raw identity | Composition | Manuscript role |
| --- | --- | --- | --- | --- |
| highly_novel | 16 | 0.802–0.835 | 16 HLA-DQ heterodimers | Reported in the Supplementary Information only; a stricter test than the headline stratum |
| strict_stratum | 39 | 0.862–0.896 | 34 HLA-C + 5 HLA-DQ | Underlies the headline strict stratum; the scored PA-med 0.765 set is the $\geq 5$-positive/$\geq 5$-negative-qualifying subset of all $< 0.90$ alleles (see caption) |
| marginal_stratum | 29 | 0.901–0.929 | 17 HLA-B + 7 HLA-DQ + 5 HLA-C | Reported as the Table 1 near-twin substratum (PA-med 0.661) |
| bridge_zone | 0 | $\geq 0.93$ | — | Empty by construction; no test allele remains within the bridge zone after the purge |
| *Most-at-risk strict-stratum alleles (max identity 0.896, nearest to bridge zone):* |  |  |  |  |
| HLA-C*03:04, HLA-C*12:02, HLA-C*08:01, HLA-C*12:03, HLA-C*16:01 (all HLA-C with nearest training neighbor an HLA-B) |  |  |  |  |
| *Most-distant highly-novel alleles (max identity 0.802):* |  |  |  |  |
| HLA-DQA1*01:02-DQB1*05:01, HLA-DQA1*01:05-DQB1*05:01, HLA-DQA1*01:05-DQB1*05:02 (HLA-DQ heterodimers at 0.802 to nearest training neighbor) |  |  |  |  |

**Table 5:** V15b pseudosequence ablation: 182-residue G-domain versus 34-residue NetMHC-pan/NetMHCIIpan pseudosequence. Both ensembles trained with the production recipe (3 seeds; frozen ESM-2 650M backbone; dropout self-attention head; matched per-seed *d*_model_ 512/256/256; same V15b bridge-purged corpus and same train/val/test splits) and scored on the 39-allele cluster-AND-sequence-disjoint experimental (eluted-ligand-dominated) stratum used for the manuscript’s lower-bound PA-med of 0.765 (Table 1). The ablation’s single delta from the open ensemble is the MHC representation: a 182-residue G-domain (open) versus a 34-residue pseudosequence (this ablation) per allele. Class I pseudosequence positions are the 34 NetMHCpan-4.0 contact positions^21^ extracted from the V15 platform sequence; Class II pseudosequences are the canonical NetMHCIIpan-2016 dictionary^27^, distributed bit-for-bit identical by both DeepMHCII ^7^ and ConBoTNet^8^.

| Stratum | MHC representation | PA-med | 95% CI |
| --- | --- | --- | --- |
| *39-allele cluster-AND-sequence-disjoint experimental, eluted-ligand-dominated (manuscript headline stratum)* |  |  |  |
| All 39 alleles | 182-residue G-domain (open) | **0.7649** | [0.691, 0.797] |
| All 39 alleles | 34-residue pseudosequence | 0.7337 | [0.638, 0.762] |
| $\Delta$ (pseudoseq – G-domain), paired Wilcoxon $p = 0.013$ | | **–0.031** | — |
| *Class breakdown (same stratum)* |  |  |  |
| 23 Class I (HLA-C at this stratum) | 182-res G-domain | 0.7680 | — |
|  | 34-res pseudoseq | 0.7101 | — |
| $\Delta$ | | –0.058 | — |
| 16 Class II (HLA-DQ/DPA-DPB) | 182-res G-domain | 0.7090 | — |
|  | 34-res pseudoseq | 0.7524 | — |
| $\Delta$ | | +0.044 | — |
| *Per-bucket $\Delta$ on broad V15b experimental test (all clean alleles)\** |  |  |  |
| both_novel ($n_a = 27$) | cluster-AND-sequence-disjoint set-<br>ting | –0.036 | G-domain higher |
| novel_peptide ($n_a = 114$) | known alleles, new peptides | +0.006 | Comparable |
| novel_allele ($n_a = 26$) | new alleles, known peptides | +0.033 | Pseudosequence higher |

Trimming the 4 HLA-C alleles with $n < 30$ test rows ( $|\Delta| > 0.5$ each, driven by perfect G-domain separation of very small samples) gives 35 alleles with $\Delta = -0.007$ (essentially tied). The G-domain’s advantage on the all-39 stratum therefore concentrates in small-sample rare-allele edge cases. \*Per-bucket $\Delta$ is the median-over-alleles of per-allele AUROC (PA-med), pseudoseq – G-domain. Source artifacts: `outputs/v15_pseudoseq_ablation/{pseudoseq_ablation_verdict.json, per_allele_delta_39.csv, final_analysis.json}`; filter chain verbatim from `scripts/laptop_peptide_only_baseline_v38.py` (the verified producer of the open-ensemble PA-med 0.7649). Audit trail at `docs/manuscript/audit.md` gA72b.

**Table 6:**
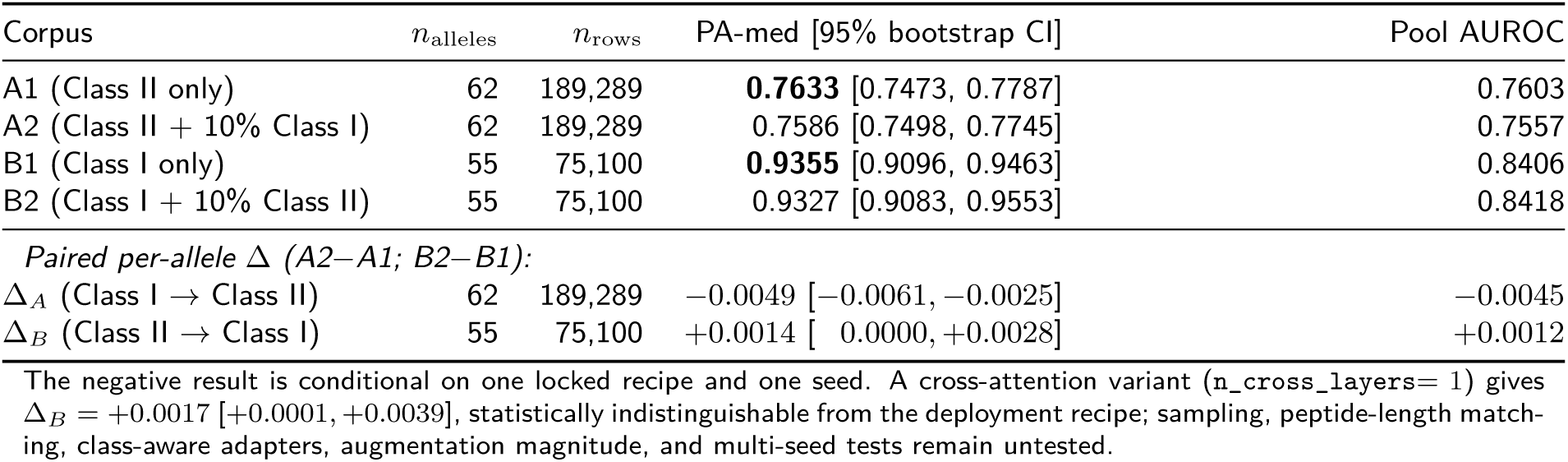
Cross-class augmentation ablation: shared-parameter head shares capacity but not learned features across MHC classes. Four single-seed deployment-recipe runs (frozen ESM-2 650M backbone, 3.9M-parameter shared head, 6 epochs, lr = 10^−4^, focal-*α* = 0.43). Targets A1 (Class II baseline), A2 (Class II + 10% Class I), B1 (Class I baseline), B2 (Class I + 10% Class II). All metrics computed on the V15 1% held-out val partition restricted to the target class via a class-strict allele-name filter (Class II target = HLA-D*; Class I target = HLA-[ABC]*), which drops 9,832 mis-tagged rows of Class I alleles that 11-source aggregation tags mhc_class = II in some sources (per Table 1 footnote). The two paired-Δ rows (62 Class II, 55 Class I alleles) are the authoritative numbers. Run-level artifacts on Zenodo (Data availability).

| Corpus | $n_{\text{alleles}}$ | $n_{\text{rows}}$ | PA-med [95% bootstrap CI] | Pool AUROC |
| --- | --- | --- | --- | --- |
| A1 (Class II only) | 62 | 189,289 | **0.7633** [0.7473, 0.7787] | 0.7603 |
| A2 (Class II + 10% Class I) | 62 | 189,289 | 0.7586 [0.7498, 0.7745] | 0.7557 |
| B1 (Class I only) | 55 | 75,100 | **0.9355** [0.9096, 0.9463] | 0.8406 |
| B2 (Class I + 10% Class II) | 55 | 75,100 | 0.9327 [0.9083, 0.9553] | 0.8418 |
| *Paired per-allele $\Delta$ (A2–A1; B2–B1):* |  |  |  |  |
| $\Delta_A$ (Class I → Class II) | 62 | 189,289 | –0.0049 [–0.0061, –0.0025] | –0.0045 |
| $\Delta_B$ (Class II → Class I) | 55 | 75,100 | +0.0014 [ 0.0000, +0.0028] | +0.0012 |

The negative result is conditional on one locked recipe and one seed. A cross-attention variant (`n_cross_layers=1`) gives $\Delta_B = +0.0017$ [ $+0.0001, +0.0039$ ], statistically indistinguishable from the deployment recipe; sampling, peptide-length matching, class-aware adapters, augmentation magnitude, and multi-seed tests remain untested.

**Table 7:** Per-row decoy distinguishability of the open ensemble. For 26 test alleles with adequate experimental-negative and generated-decoy counts in the 1% open-ensemble val partition, AUROC for distinguishing experimental negatives (label = 0, real non-binders) from generated decoys (label = 0, synthetic non-binders). Both classes share binding_label = 0, so AUROC 0.5 means the model does not separate them; AUROC *>* 0.55 means the model has learned a decoy-generation artifact; AUROC *>* 0.7 would seriously compromise headline interpretation. Median across alleles is at chance (0.516); zero alleles exceed 0.70; the corpus-level Δ between full and experimental-only val partitions is −0.031, so including decoys *lowers* the headline rather than inflating it.

| Five lowest AUROC |  | Five near-median AUROC |  | Five highest AUROC |  |
| --- | --- | --- | --- | --- | --- |
| Allele | AUROC | Allele | AUROC | Allele | AUROC |
| HLA-DRB1*13:02 | 0.020 | HLA-DRB3*01:01 | 0.488 | HLA-B*44:02 | 0.618 |
| HLA-DRB1*15:01 | 0.085 | HLA-A*02:01 | 0.511 | DQA1*02:01-DQB1*02:02 | 0.625 |
| HLA-DRB1*01:01 | 0.097 | HLA-A*03:01 | 0.522 | HLA-B*27:05 | 0.644 |
| HLA-A*02:05 | 0.135 | HLA-A*23:01 | 0.530 | HLA-B*40:01 | 0.646 |
| HLA-DRB1*07:01 | 0.160 | DQA1*03:03-DQB1*02:02 | 0.534 | HLA-A*24:02 | 0.678 |

*Aggregate over all 26 alleles:*
Median AUROC **0.516** (centred at chance) · IQR [0.260, 0.582] · range [0.020, 0.678]
Fraction > 0.55: 30.8% (mild signal on a minority) · Fraction > 0.60: 19.2% · **Fraction > 0.70: 0%**
Pool-level $\Delta$ (full minus experimental-only val): –0.0312 (decoys slightly *lower* the headline)
Low-AUROC alleles (HLA-DRB1\*13:02 etc.) indicate that, for the open ensemble, generated decoys receive more binder-like scores than experimental negatives on those alleles rather than the reverse, again consistent with absence of a decoy-recognition shortcut. Full 26-row JSON deposited on Zenodo (`outputs/v37_decoy_audit/audit.json`).

**Table 8:** Cluster-novel check across three disjoint held-cluster compositions. Each fold trains an identical production-recipe ensemble on a separate corpus where one set of held alleles plus all bridges at ≥0.90 raw G-domain identity to those alleles have been purged. Fold compositions are not directly comparable (different gene families held to different completeness), so per-fold PA-med is reported rather than a single cross-fold mean. The headline lower bound in Table 1 (PA-med 0.765 on 39 alleles) uses a hand-curated 30-allele within-family bridge drop; this design applies an automatic, uniformly stricter 90%-identity bridge purge.

| fold | seeds | held cluster (representative) | $n_{\text{CLEAN}}$ | ba_exp PA-med | per-gene story |
| --- | --- | --- | --- | --- | --- |
| 1 | 2 | HLA-C*03:04 + DPA1*01:03-DPB1*02:01<br>+ DQA1*02:01-DQB1*06:04 + E*01:03 | 30 | 0.586 | HLA-B (7) 0.526; HLA-C (13) 0.575; HLA-DP_het (10) 0.665 |
| 2 | 1 | HLA-A*23:01 + DQA1*01:02-DQB1*06:02 | 11 | 0.550 | HLA-A (4) 0.499; DQ_het (7) 0.550 |
| 3 | 1 | HLA-B*57:01 + A*01:01 + DQA1*01:02-DQB1*02:02 + DQA1*03:02-DQB1*04:01 | 15 | 0.566 | HLA-B (9) 0.693; HLA-A (2) 0.418; DQA1*03 het (4) 0.310 |

**Table 9:**
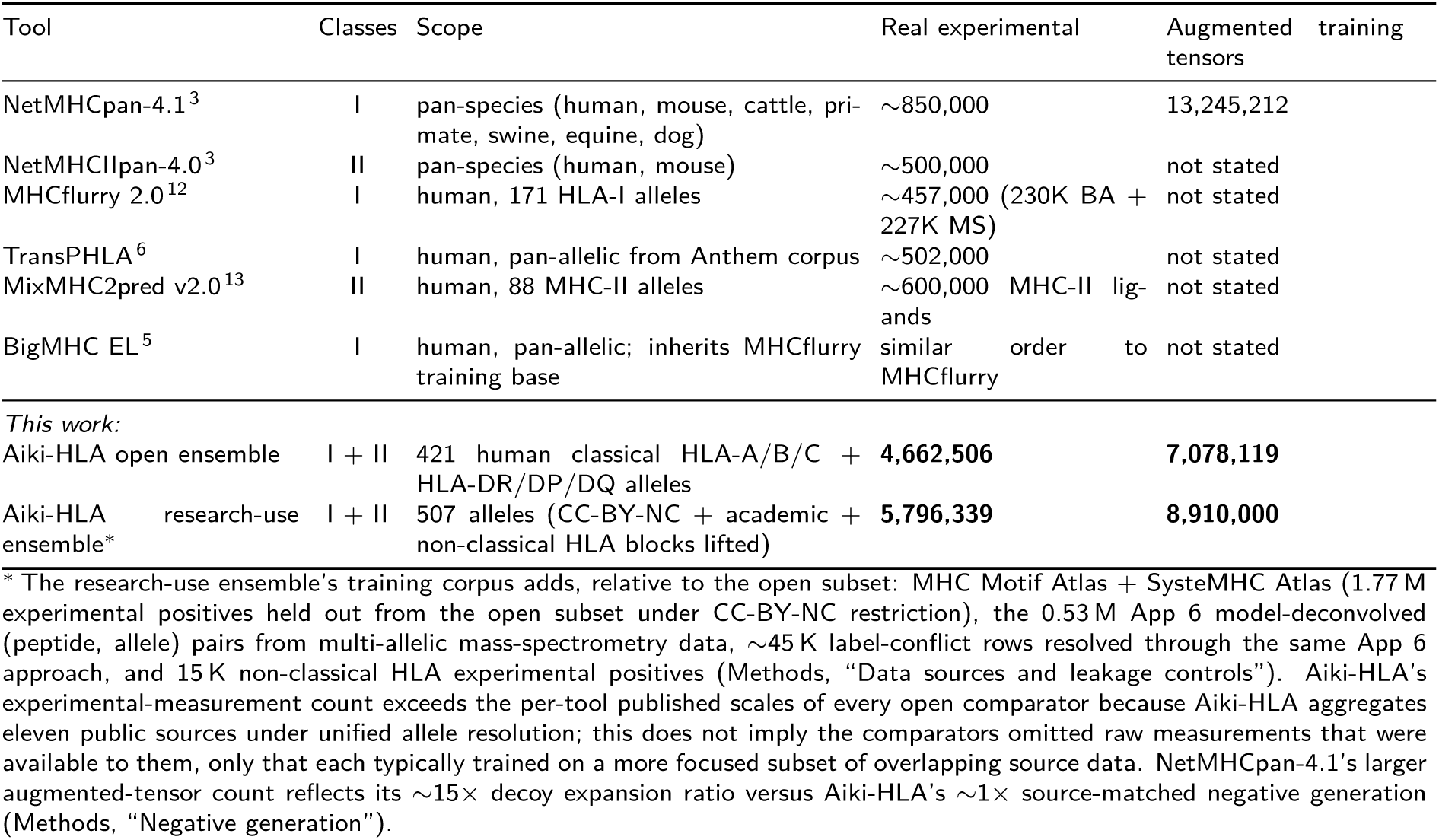
Training-corpus scale comparison across open pMHC predictors. “Experimental measurements” refers to the real peptide-MHC observations (binding-affinity assays + mass-spectrometry eluted ligands) reported by each tool’s primary publication, before each tool’s own decoy augmentation. “Augmented training tensors” refers to the total data points each tool optimizes against, including each tool’s generated decoys. The two columns are not strictly comparable because each tool augments differently (NetMHCpan-4.1 uses 15 decoy expansion; Aiki-HLA uses 1 ); the experimental-measurement column is the cleaner comparison of source-data scale. Aiki-HLA’s experimental count is larger than each single comparator because Aiki-HLA aggregates eleven public sources under unified allele resolution rather than because comparators left raw measurements unused; most published tools train on a focused subset of those same sources.

| Tool | Classes | Scope | Real experimental | Augmented training tensors |
| --- | --- | --- | --- | --- |
| NetMHCpan-4.1<sup>3</sup> | I | pan-species (human, mouse, cattle, primate, swine, equine, dog) | $\sim 850,000$ | 13,245,212 |
| NetMHCIIpan-4.0<sup>3</sup> | II | pan-species (human, mouse) | $\sim 500,000$ | not stated |
| MHCflurry 2.0<sup>12</sup> | I | human, 171 HLA-I alleles | $\sim 457,000$ (230K BA + 227K MS) | not stated |
| TransPHLA<sup>6</sup> | I | human, pan-allelic from Anthem corpus | $\sim 502,000$ | not stated |
| MixMHC2pred v2.0<sup>13</sup> | II | human, 88 MHC-II alleles | $\sim 600,000$ MHC-II ligands | not stated |
| BigMHC EL<sup>5</sup> | I | human, pan-allelic; inherits MHCflurry training base | similar order to MHCflurry | not stated |
| *This work:* |  |  |  |  |
| Aiki-HLA open ensemble | I + II | 421 human classical HLA-A/B/C + HLA-DR/DP/DQ alleles | **4,662,506** | **7,078,119** |
| Aiki-HLA research-use ensemble* | I + II | 507 alleles (CC-BY-NC + academic + non-classical HLA blocks lifted) | **5,796,339** | **8,910,000** |

\* The research-use ensemble’s training corpus adds, relative to the open subset: MHC Motif Atlas + SysteMHC Atlas (1.77 M experimental positives held out from the open subset under CC-BY-NC restriction), the 0.53 M App 6 model-deconvolved (peptide, allele) pairs from multi-allelic mass-spectrometry data, $\sim 45$ K label-conflict rows resolved through the same App 6 approach, and 15 K non-classical HLA experimental positives (Methods, “Data sources and leakage controls”). Aiki-HLA’s experimental-measurement count exceeds the per-tool published scales of every open comparator because Aiki-HLA aggregates eleven public sources under unified allele resolution; this does not imply the comparators omitted raw measurements that were available to them, only that each typically trained on a more focused subset of overlapping source data. NetMHCpan-4.1’s larger augmented-tensor count reflects its $\sim 15\times$ decoy expansion ratio versus Aiki-HLA’s $\sim 1\times$ source-matched negative generation (Methods, “Negative generation”).

